# Dynamically shifting from compositional to conjunctive brain representations supports cognitive task learning

**DOI:** 10.1101/2023.06.27.546751

**Authors:** Ravi D. Mill, Michael W. Cole

## Abstract

During cognitive task learning, neural representations must be rapidly constructed for novel task performance, then optimized for robust practiced task performance. How the geometry of neural representations changes to enable this transition from novel to practiced performance remains unknown. We hypothesized that practice involves a shift from compositional representations (task-general activity patterns that can be flexibly reused across tasks) to conjunctive representations (task-specific activity patterns specialized for the current task). Functional MRI during learning of multiple complex tasks substantiated this dynamic shift from compositional to conjunctive representations, which was associated with reduced cross-task interference (via pattern separation) and behavioral improvement. Further, we found that conjunctions originated in subcortex (hippocampus and cerebellum) and slowly spread to cortex, extending multiple memory systems theories to encompass cognitive task learning. The strengthening of conjunctive representations hence serves as a computational signature of learning, reflecting cortical-subcortical dynamics that optimize task representations in the human brain.

**Highlights:**

- Learning shifts multi-task representations from compositional to conjunctive formats
- Cortical conjunctions uniquely associate with improved behavior and pattern separation
- These conjunctions strengthen over separated learning events and index switch costs
- Subcortical regions are critical for cross-region binding of task rule information

## Introduction

Learning cognitive tasks is a common feature of everyday life, yet the neural basis for this essential human skill remains unknown. Studies have consistently reported behavioral improvement as tasks are performed from initial novel presentation through extensive practice, which has been linked to a progression from controlled to automatic processing (Schneider & Chein, 2003; Schneider & Shiffrin, 1977). fMRI has alluded to neural changes accompanying this progression, in the form of widespread decreases in the magnitude and extent of regional activations over practice, with some spatially limited increases (Chein & Schneider, 2005; Hampshire et al., 2019; Kami et al., 1995; Poldrack, 1998; Ruge & Wolfensteller, 2010). The activation decreases are most prominent in cognitive control-related regions (Bassett et al., 2015; Cole et al., 2010; Mohr et al., 2016). However, a precise computational account for how neural representations change over learning remains elusive.

In the present report, we assimilated concepts from the language, long-term memory, context-dependent decision-making and cognitive control neuroscientific literatures to develop novel hypotheses about the neural basis of task learning. These hypotheses are elaborated below, and summarized here: i) novel task performance is reliant on task-general compositional brain representations, ii) practiced task performance is reliant on task-specialized conjunctive brain representations, iii) these representational dynamics are differentially evoked in cortex versus subcortex (constituting a division of labor between these systems), and iv) progressive strengthening of conjunctions specifically in cortex relates to signatures of effective task practice.

Our first hypothesis was informed by early language theories (Chomsky, 1957; Montague, 1970) and ongoing research involving artificial neural networks (Lake et al., 2015; McClelland & Rumelhart, 1985; Yang et al., 2019), which position compositionality as an important representational principle. Compositions provide a format for representing abstract information that can be retrieved and recombined based on task context, theoretically allowing for flexible thought and behavior (Behrens et al., 2018; Cole et al., 2011; Collins & Frank, 2013; Reverberi et al., 2012). Interest in compositionality has been accelerated by observed shortcomings of artificial neural network models in performing rapid instructed task learning (Cole et al., 2013) - a human ability allowing for competent task performance with little to no practice. Longstanding artificial neural network deficits in transferring prior learning in “rapid” task contexts (e.g. zero-shot transfer; Lake et al., 2017; Palatucci et al., 2009) have recently been ameliorated by models embedded with compositional principles (Ito et al., 2022; Yang et al., 2019). Extending these findings to human cognitive task learning, we predicted that compositional representations in the brain play a critical role in processing novel tasks. Critically, the degree to which compositional representations are evoked in empirical brain data (versus simulated data used for artificial neural network modeling), and how this might vary over the course of extended task learning, remains uncertain.

Whilst compositional representations theoretically allow for task competency in novel contexts, compositionality alone is unlikely to account for established effects of practice in improving behavioral performance towards task proficiency. This informed our second hypothesis, which frames a complementary role for conjunctive representations as the neural substrate of task practice. In long-term memory research, nonlinear conjunctions of sensory features have long been theorized as a key computation, under varying terminology: chunking (Wickelgren, 1979), binding (Ranganath, 2010; Shimamura & Wickens, 2009; Yonelinas et al., 2019), relational representation (Eichenbaum, 2013), configural (versus elemental) association (Marr, 1971; Sutherland & Rudy, 1989) and object-based (versus feature-based) representation (Radulescu et al., 2021). Extending these ideas to the domain of cognitive task learning, we predicted that conjunctions of task rule elements would strengthen over extended learning, and uniquely associate with signatures of effective practice (reduced representational interference and improved behavior). This would offer a contrasting trajectory to compositional representations, which would be more strongly evoked early in learning, and then progressively weaken.

Finally, we also predicted that these representational dynamics would be differentially evoked across cortical and subcortical systems. This was founded on seminal neuropsychological studies of long-term memory demonstrating that medial temporal lobe (MTL) damage impairs certain forms of memory, such as episodic memory (Scoville & Milner, 1957), whilst preserving others, such as procedural skill learning (Brooks & Baddeley, 1976; Corkin, 1984; Eichenbaum, 2013; Squire, 1992). This pattern of deficits highlighted that regions beyond MTL also subserve learning and memory, which was formalized in many multiple memory systems frameworks (Hirsh, 1980; Manns & Eichenbaum, 2006; Nadel et al., 2000; O’Reilly & Rudy, 2001; Sherry & Schacter, 1987; Simons & Spiers, 2003; Sutherland & Rudy, 1989).

Critical amongst these was complementary learning systems theory (Kumaran et al., 2016; McClelland et al., 1995; O’Reilly et al., 2014; O’Reilly & Rudy, 2001), which posited differentiable roles for MTL and cortical regions in learning. MTL was theorized to perform fast learning (via fast conjunction formation) whereas cortex performs slow learning (via slow conjunction formation). These distinct roles in the service of a common learning function was characterized as a “division of labor” between cortical and subcortical systems. Formalizing this division of labor in artificial neural network models accurately simulated memory deficits in animal lesioning studies (O’Reilly & Rudy, 2001). Extending to cognitive task learning, we predicted that repeated task practice is underpinned by strengthening of conjunctive representations in cortex specifically. This follows from the alignment of the theorized slow learning function of cortex with the slow timescale of practice-related behavioral improvement. Critically, to our knowledge, the predictions of complementary learning systems theory have not been tested in empirical human brain data - in the domain of task learning, but also generally.

Furthermore, complementary learning systems theory places predominant emphasis on the role of conjunctive representations in extended learning, thereby neglecting the reciprocal importance of compositions under task novelty. This contrasts with the inverse pattern of emphasis in recent artificial neural network models - promoting the importance of compositions (as a substrate of abstraction) and neglecting conjunctions (as a substrate of specialization). We offer an integrative theoretical account in Fig 1A, in which cortical-subcortical dynamics underpin a shift from relying on compositional representations when tasks are novel, to relying on conjunctive representations with task practice. We conceptualize subcortical involvement in learning as not exclusive to the MTL (per complementary learning systems), but also strongly involving the cerebellum, given its prior linkages with task learning (Beukema & Verstynen, 2018; Ferrari et al., 2018), behavioral automaticity (Chein & Schneider, 2005; Lang & Bastian, 2002; Nixon & Passingham, 2000), and flexibility generally (J.-H. Gao et al., 1996; King et al., 2019; Nakai & Nishimoto, 2022). Fig 1B schematizes how cortical conjunctive strengthening might drive signatures of effective task practice - reduced representational interference and improved behavior.

**Fig 1.**
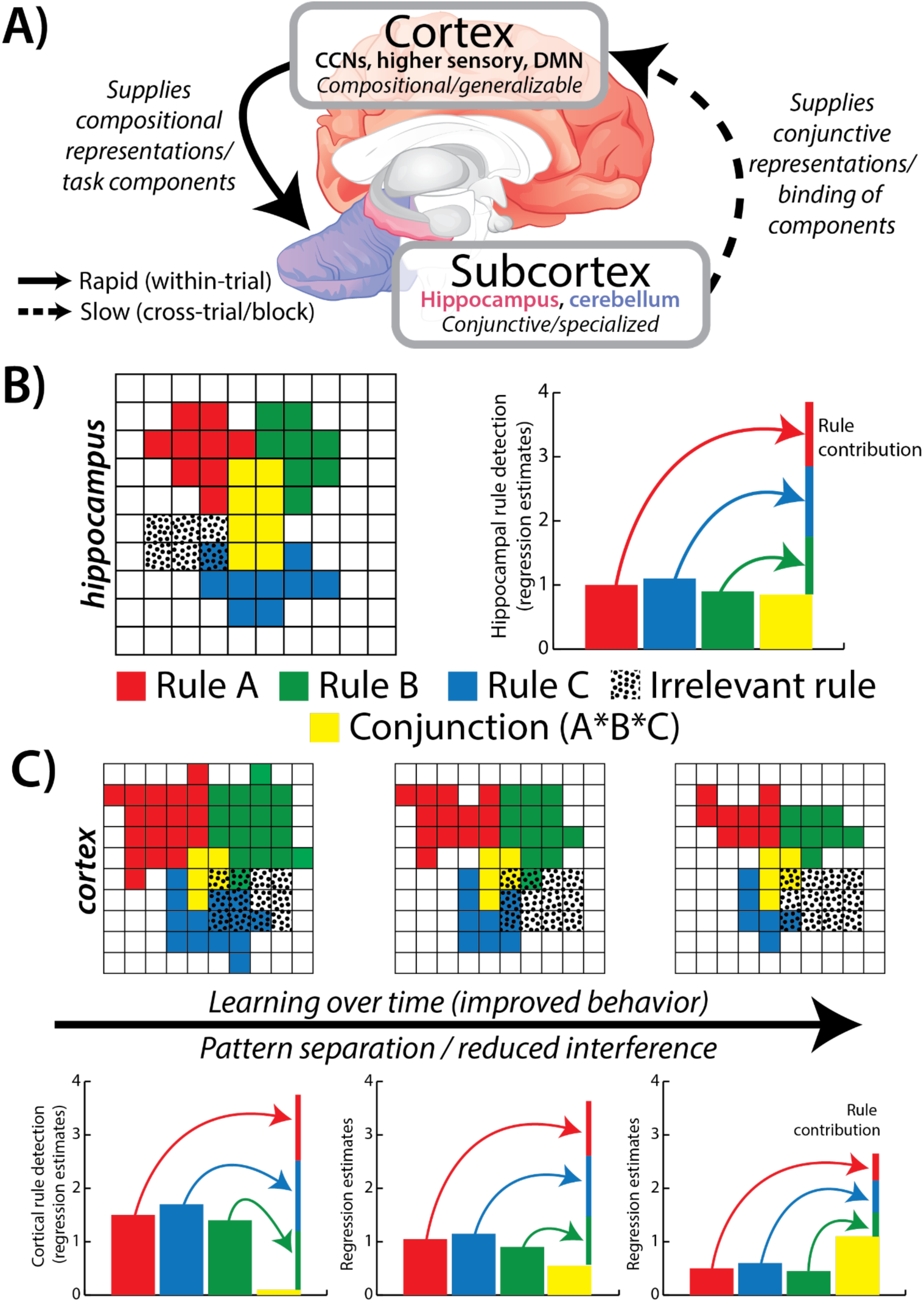
Hypothesized cortical-subcortical dynamics underlying changes to neural representational geometry over task learning. **A)** Depiction in neural space. Early in learning, compositional representations of task rules are retrieved and reinstated in cortical networks (including cognitive control networks, CCNs, higher sensory networks and the default mode network, DMN), facilitating transfer of relevant pre-experiment knowledge to the current context (here, performance of C-PRO2 paradigm tasks; see Fig 2A for design). These compositional representations are then rapidly bound by subcortex (hippocampus and cerebellum prominently) into more specialized conjunctive representations, and over practice propagated back to cortex, where they gradually strengthen over the compositional representations to facilitate behavioral improvement. **B)** Depiction in representational space: hypothesized spatial activity pattern evoked in hippocampus early in learning. Complementary depictions of the same representational profile are provided: the left panel presents the hypothetical within-region activation pattern (each square representing activation of within-region voxels), and the right panel presents hypothetical results from applying our machine learning approach to quantify rule representation strength in this region (i.e. regression weights from fitting linear compositional and non-linear conjunctive rule templates to the evoked activation pattern; see Fig 4A for details). Early in learning, hippocampus binds compositional task rule representations into non-linear conjunctions, which promotes encoding of a more distinct, specialized representation of the current task. One computational benefit of such conjunctive coding is to allow for a pattern-separated representation of the task, which is more resistant to interference from irrelevant rules (depicted as reduced overlap of task-relevant rules with an irrelevant rule’s dotted pattern). **C)** Equivalent activation (top panel) and regression space (bottom panel) depictions for a cortical region over learning. As tasks are repeatedly practiced, conjunctive codes increasingly dominate in cortical regions, with reliance on compositional codes reduced in parallel. The computational benefit of these representation dynamics is to promote task pattern separation, reducing interference with neural encoding of other tasks, and with a particular impact (compared to subcortex) in improving behavioral performance with extended practice.

In devising an analytic framework to test the schematized learning principles in empirical human brain data, we were inspired by recent innovations in the context-dependent decision-making literature. Machine learning and multivariate pattern analysis approaches have been applied to functional brain data to quantify the degree to which neural representations evoked during decision-making are context-specific (conjunctive) or context-general (compositional) (Badre et al., 2021; Bernardi et al., 2020; Dang et al., 2021; Flesch et al., 2022; Fusi et al., 2016; Ito et al., 2020; Kikumoto & Mayr, 2020; Mack et al., 2020; Mante et al., 2013; Rigotti et al., 2013). Whereas these representational geometry approaches have been primarily applied at the within-trial timescale, here we applied similar methods over the longer timescale of extended task learning.

We tested our hypotheses using fMRI recordings in healthy adult human participants as they performed multiple complex cognitive tasks from initial novel presentation through repeated practice. We developed a new version of the concrete permuted rule operations paradigm (C-PRO2), which preserved the compositional (each task is created by permuting sensory, logic and motor rule types) and multi-task (allowing for more generalizable inferences than single-task approaches) nature of previous versions (Cole et al., 2010; Cole, Reynolds, et al., 2013; Ito et al., 2017), but now with more complex, naturalistic and diverse features (see Methods and Fig 2A). Collectively, this combination of theoretical, analytic and task design innovations allowed us to empirically substantiate a neurocomputational signature of task learning (Fig 1), in which cortical-subcortical dynamics underpin a transition from compositional to conjunctive representations with practice.

**Fig 2.**
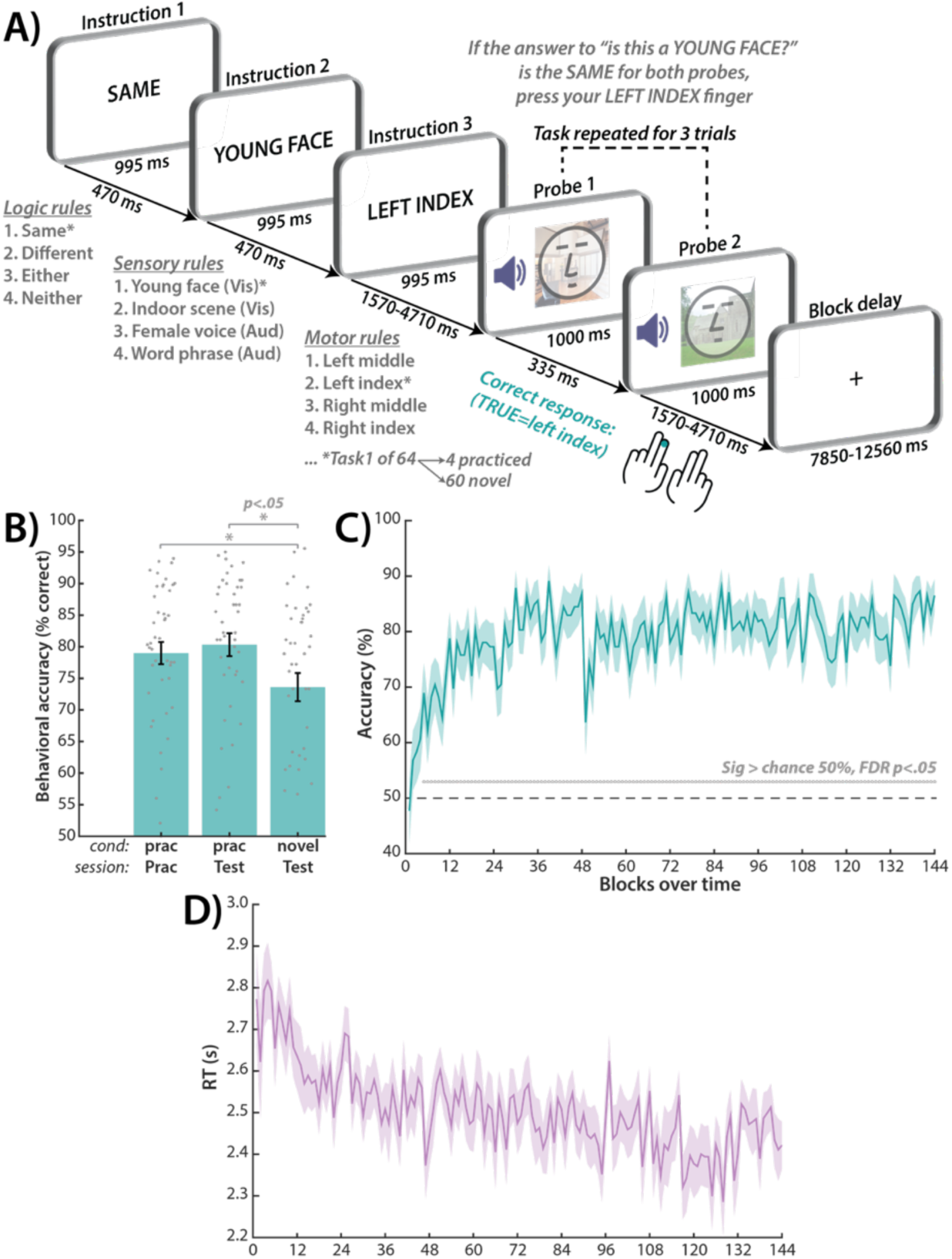
C-PRO2 task design and observed behavioral learning effects. **A)** Format for one block of the C-PRO2 task paradigm (see Methods for details), with an example trial and accompanying correct response. This block format was preserved for all 64 tasks, across practiced/novel task conditions and Practice/Test sessions. A task block began with three instruction screens (one each for logic, sensory and motor rule types), which conveyed the rules to be applied for the ensuing set of three trials. Each trial concurrently presented composite visual (dimensions: young or old face, indoor or outdoor scene) and composite auditory (dimensions: female or male voice, word or non-word content) stimuli over two sequential probes. Subjects were required to integrate the task rule information to process the incoming stimulus information across the probes, and make their true/false categorization responses. **B)** Session-averaged accuracy across key task conditions: practiced tasks in the first Practice session, practiced tasks in the second Test session, novel tasks in the Test session. Subjects were reliably above chance performance, and practiced tasks were better performed than novel ones (asterisks denote significance of paired t-tests, p<.05). Error bars reflect standard error of the mean, with subject datapoints overlaid. **C)** Accuracy over task blocks in the Practice phase, showing expected learning effects: rise in accuracy over time (significantly above-chance by block 5 via one-sample t-test; significance denoted by gray dots at FDR p<.05) that plateaued around 30 blocks. **D)** Reaction time (RT) over task blocks in the Practice phase, also showing improvement in performance over learning (i.e. quickening of RT). Patches in C) and D) reflect standard error of the mean.

## Results

### Behavioral evidence of task learning in novel C-PRO2 paradigm

We first verified that task learning had taken place as expected in our new C-PRO2 paradigm (see Methods and Fig 2A for task details). Fig 2B plots the block-averaged accuracy across key conditions: practiced tasks presented in the first Practice session (4 tasks repeated 36 blocks each), practiced tasks in the second Test session (same 4 tasks repeated for a further 15 blocks each), and novel tasks in the Test session (60 tasks presented once). Subjects were reliably above chance accuracy (cross-task average=77.6%) in all three conditions (one-sample t-tests against 50% chance, p<.05), confirming that participants understood the paradigm. The cross-condition profile demonstrated that practice improved behavior: practiced tasks in both the Practice (mean=79.0%) and Test (80.3%) sessions were more accurately performed than novel (73.6%) tasks (practiced task in Practice session > novel task in Test session, t=2.90, p=.006; practiced task in Test session > novel task in Test session, t=5.14, p<.001). Accuracy for practiced tasks in the Test session was numerically albeit non-significantly higher than practiced tasks in the Practice session (t=1.10, p=.276), and there were no reliable between-condition differences in RT (paired t-tests, all p>.450). Further evidence of learning came from timecourse analyses of performance accuracy (Fig 2C) and reaction time (RT; Fig 2D) over Practice session blocks. In both cases, behavior improved (accuracy increased and RT quickened) with repeated practice (paired t-test for last block > first block improvement, accuracy t=6.27, p<.001; RT t=2.47, p=0.018). These results confirmed that the key task learning manipulation was successful.

### Evidence of task pattern separation: neural task similarity decreases with practice to reduce interference

Our fMRI analyses generally adopted multivariate pattern analysis techniques. We first used a classification approach (4-way decoding of practiced task identity) as a sanity check to verify that task information was reliably decodable from multivariate neural activations evoked by our paradigm (Fig S1).

Subsequent analyses targeted the critical question of how the geometry of these neural representations changed over learning. We tested whether representational similarity in the brain decreased with repeated task practice, consistent with pattern separation learning processes that reduce interference. We hypothesized this as a key signature of practice (Fig 1), which we later link to more refined representational dynamics (Fig 7). Here, our goal was to preliminarily establish pattern separation in the domain of cognitive task learning, versus more typically studied spatial memory and object recognition domains.

Our method of estimating neural task similarity is depicted in Fig 3A. We analyzed the resulting regional similarity timecourses in two complementary ways, which, coupled with our rigorous fMRI preprocessing steps, ruled out potential contamination by temporal artifacts (see Methods). Firstly, we estimated the dynamic trajectory (regression slope over time) of the practiced task similarity timecourses in the first Practice session. Secondly, we contrasted the block-averaged similarity of practiced versus novel tasks presented in the second Test session. Consistent with our hypothesis that task practice reduces neural task similarity (increases pattern separation), we predicted that i) regional similarity trajectories would generally decrease over time for practiced tasks in the Practice session, ii) similarity would be lower for practiced versus novel tasks in the Test session. The results find support for both predictions (see Supplementary Table for a list of significant regions and accompanying statistics), with Fig 3B depicting the conjunction of these two analyses to isolate regions most strongly linked to task pattern separation. These included medial temporal lobe regions typically associated with pattern separation (bilateral hippocampus), but also cortical and subcortical regions (bilateral cerebellum, bilateral caudate nucleus and right amygdala) not as conventionally linked.

**Fig 3.**
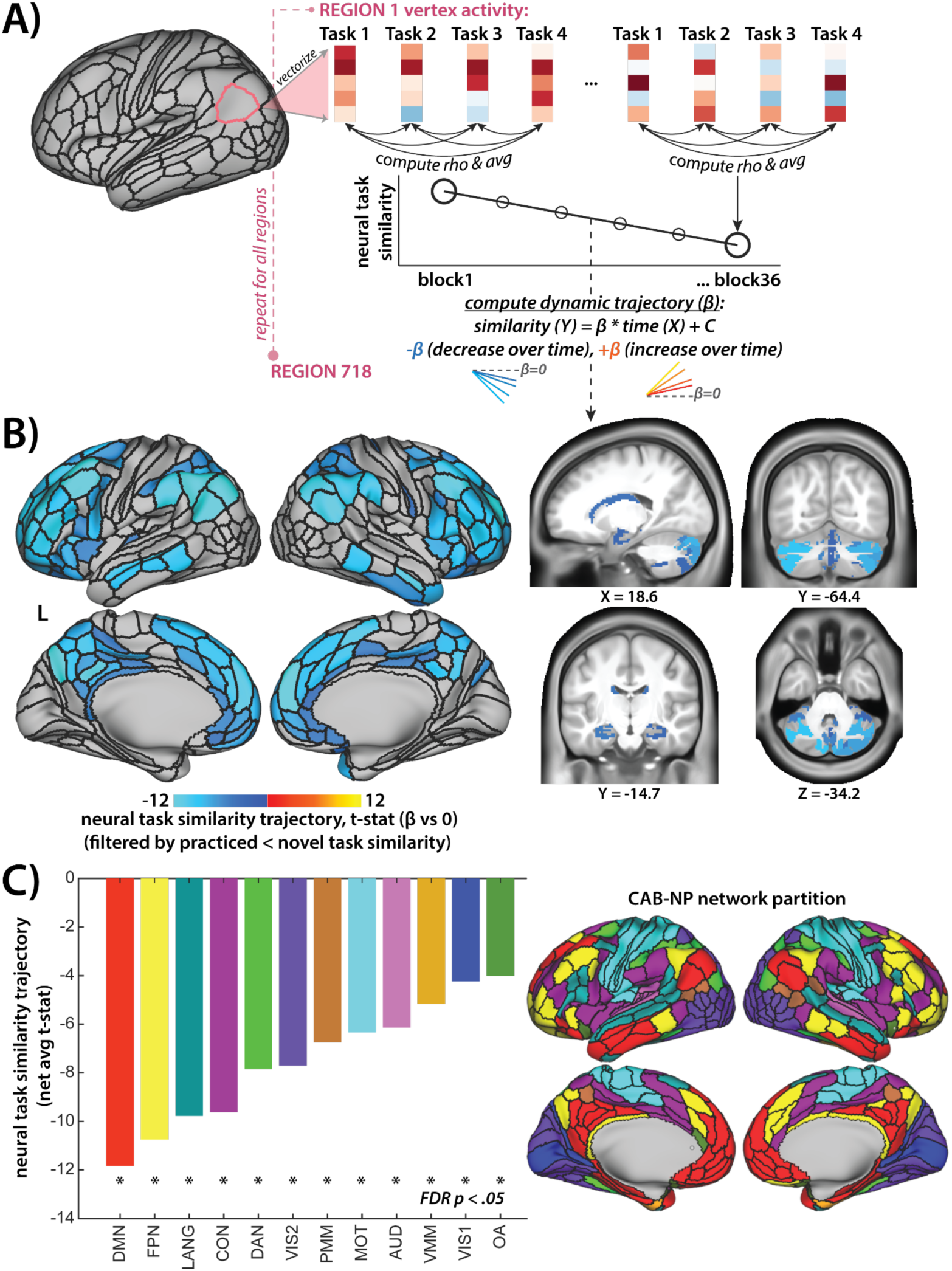
Task practice reduces neural task similarity, consistent with pattern separation processes that reduce interference. **A)** Schematic of analysis approach. For each region of interest, the similarity between task activation features (within-region vertices for cortex, and within-region voxels for subcortex) was computed using Spearman’s correlation. The average across pairwise task similarities was computed for each block, yielding a timecourse of neural task similarity over repeated practice. A regression approach then formalized the positive/negative trajectory over time in the Practice session, with this process repeated per region. **B)** Strongest pattern separation regions i.e. those showing a significant dynamic trajectory for practiced task neural similarity in the Practice session, filtered by a significant practiced < novel task similarity difference in the Test session, both at FDR p<.05. Significant cortical regions are shown in the left panel, and significant subcortical regions are shown in the right panel. Pattern separation generally increased (i.e. neural task similarity decreased) with task practice. **C)** Regional similarity trajectories in the Practice session averaged into functional networks from the CAB-NP network partition (depicted in the right panel). Networks with the strongest practice-induced increases in pattern separation were the default mode network, cognitive control networks (DAN, FPN, CON) and higher-order sensory networks (Visual 2, Language).

The Practice session regional trajectories were averaged into large-scale functional networks, based on an established network parcellation (Fig 3C). Whilst all networks showed a significant negative trajectory (FDR p<.05), this was numerically strongest in the default mode network (DMN, trajectory t=-11.83, p<.001), cognitive control networks (CON, t=-9.62, p<.001; DAN, t=-7.84, p<.001; FPN, t= −10.75, p<.001) and the higher-order sensory networks (Language, t=-9.77, p<.001; Visual 2, t=-7.71, p<.001). The supplement (Fig S2) provides results at the whole-brain level, which again revealed a negative trajectory, and further highlighted that this held generally across all 12 task rules (3 rule types * 4 levels each = 12 rules total; see Methods).

Reduction in neural task similarity over practice was also reliably associated with behavioral improvement, for both accuracy (Fig S3A) and RT (Fig S3B). Overall, the results reveal changes to neural representational geometry consistent with task pattern separation, which reduces interference with practice.

### Quantifying compositional and conjunctive rule representations during task learning reveals a dissociation between subcortex and cortex

Preceding analyses revealed that neural task representations became more distinct from each other over learning, but left unclear what specifically was changing to produce this. We hypothesized that the reduction in cross-task neural similarity over learning was driven by an increasing reliance on conjunctive (versus compositional) representations of the task rules (Fig 1).

To test this, we developed a machine learning approach to quantify the strength of compositional and conjunctive representations on a block-to-block basis (see Fig 4A and Methods for details).

**Fig 4.**
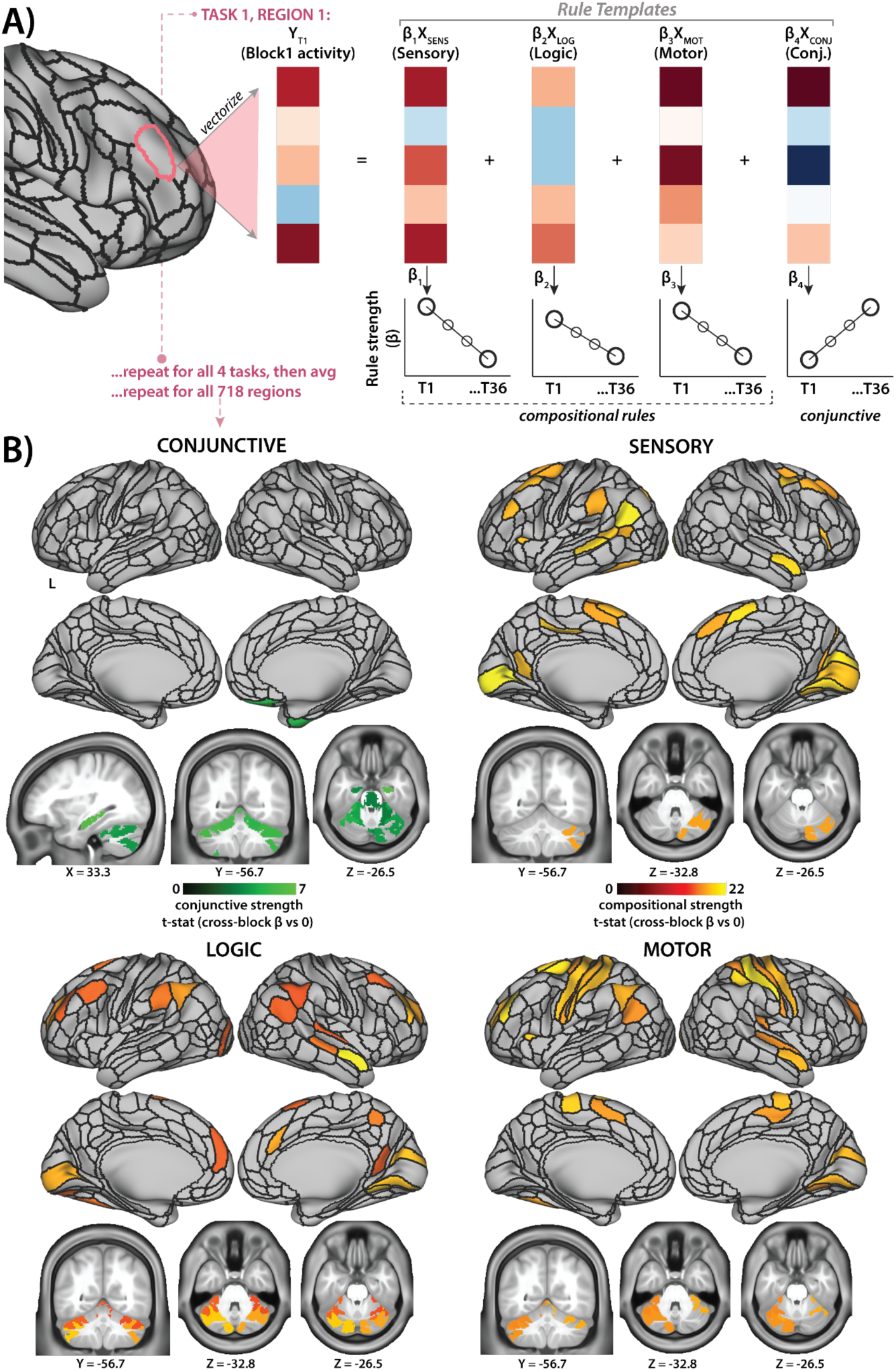
Cross-block rule strength analysis reveals differential sensitivity of subcortex and cortex to conjunctive and compositional codes. A) Schematic of our machine learning approach to detecting rule representation strengths over learning (see text for details). B) Applying the approach to quantify overall (cross-block averaged) conjunctive (top left) and compositional (clockwise from top right) rule strengths, for cortex and subcortex. To ease visualization of positive region peaks, the compositional brain plots are thresholded as the top 5% t-stats, as many regions survived FDR p<.05 correction. The conjunctive plot yielded more sparse significance and is hence depicted at the standard FDR p<.05 threshold. Conjunctive codes were strongest in subcortical regions (bilateral hippocampus, bilateral cerebellum, bilateral diencephalon, bilateral brainstem, left thalamus), with compositional codes detectable in cortex as well as subcortex. Bilateral cerebellar regions were most prominent in representing compositions (surviving the top 5% thresholding here, see Supplementary Table for statistics). Sensible peaks were obtained for each single rule type (e.g. bilateral motor cortex regions for the motor rule), validating the analytic approach.

Templates for single rule types (sensory, logic and motor) and their conjunction were estimated using held-out Test session data, and then fit simultaneously via robust regression to the regional practiced task activations, over Practice session blocks. The resulting beta coefficients quantified the strength of compositional versus conjunctive rule representations over time.

The overall (cross-block average) strength of each rule type is presented in Fig 4B. Recovery of positive region peaks for the compositional rule types, i.e. visual and auditory regions for sensory rules, lateral prefrontal regions for logic rules, and motor cortex for motor rules (FDR p<.05; see Supplementary Table for accompanying statistics), was consistent with prior literature (Mill et al., 2022; Schultz et al., 2022) and served as analysis sanity checks. The results also revealed stronger representation of conjunctions in subcortex versus cortex, with compositions strongly represented across both. This dissociable spatial profile supports our hypothesized division of labor between subcortex and cortex in fulfilling distinct learning computations.

### Rule representation dynamics reveal a neurocomputational signature of task practice: progressive replacement of compositional with conjunctive representations in cortex

To further examine dissociable roles for cortex and subcortex in task learning, we conducted dynamic analyses of rule representation strength. We hypothesized that with task practice, compositional representations in cortex would be slowly replaced with conjunctive representations (Fig 1). To ensure sufficient power, these analyses used network summaries of cortical regions (regional rule beta coefficients averaged into affiliated CAB-NP networks), and 5 subcortical regions-of-interest that showed strong pattern separation and conjunctive representation effects (see Table S1 and Fig S4).

The results are provided in Fig 5, with left columns depicting rule strength timecourses, and right columns formalizing the timecourse trajectories. Firstly, we observed significant subcortex > cortex conjunctive strength, and significant cortex > subcortex compositional strength across all blocks (all paired t-tests significant in the respective directions, for all rule types, FDR p<.05), confirming the separable representational sensitivities in the block-averaged results (Fig 4B).

**Fig 5.**
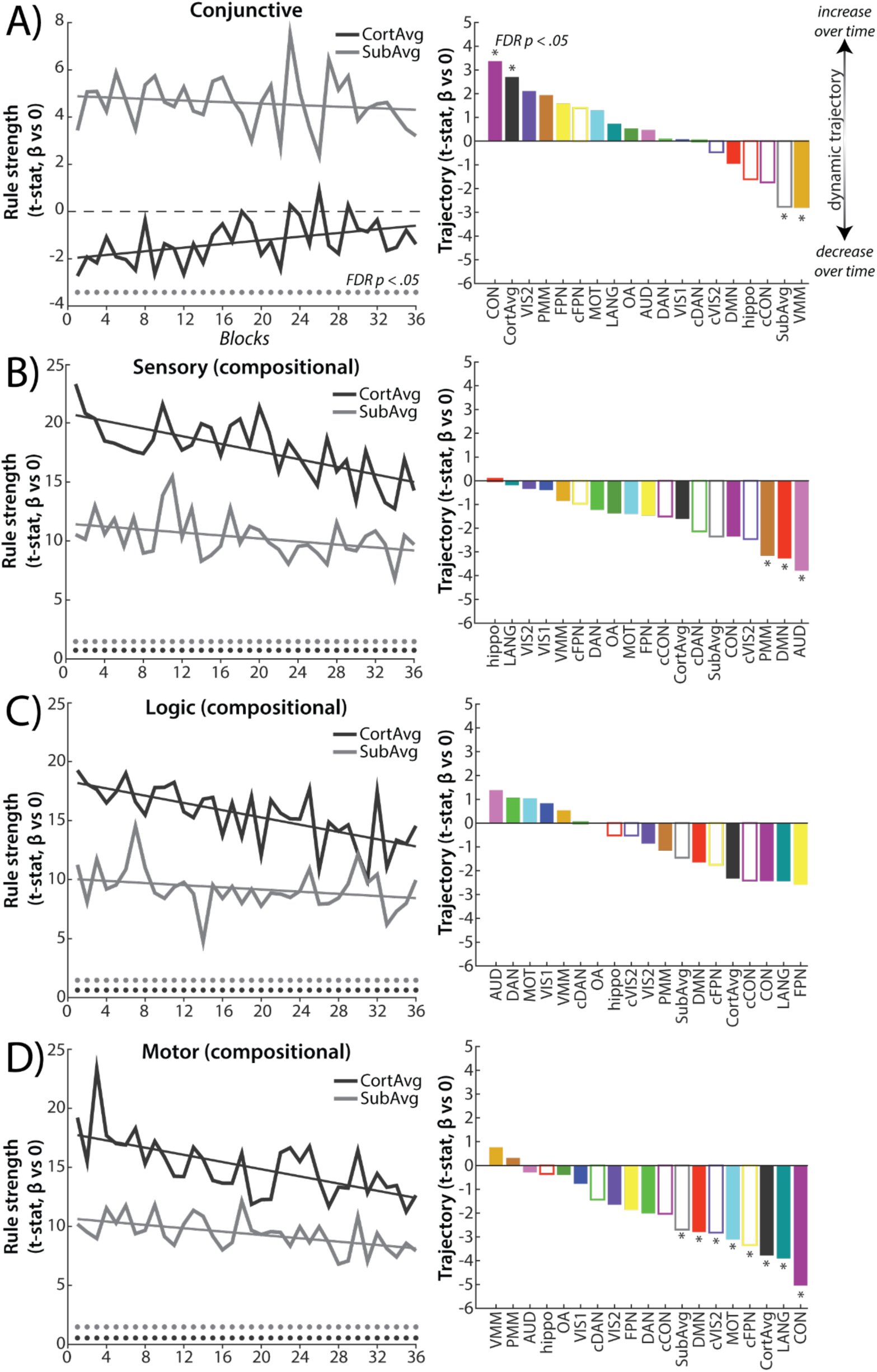
Replacement of compositional with conjunctive codes in cortex is a key computational signature of task learning. Left columns depict rule strength timecourses (t-stats for rule beta coefficients vs 0, estimated per Fig 4A) for the different rule types. To simplify presentation, depicted timecourses are the average across cortical networks (black line) and the average across subcortical regions (gray line), with linear trend lines overlaid. Dots denote significant rule strength (one-sample t-test versus 0, FDR p<.05). Right columns provide the dynamic trajectory analysis results for each cortical network and subcortical region (and their averages), which formalize the timecourse trends. Cortical networks are presented as solid bars, and subcortical regions as outlined bars. Asterisks denote significant trajectories via one-sample t-test versus 0, FDR p<.05. **A)** Dynamics of conjunctive rule representations. Whereas conjunctions were detected early in subcortex and reliably weakened over time, they were not present early in cortex but reliably strengthened over time. The dynamic trajectory analysis revealed generally increasing conjunctive strength in the cortical networks: 9/12 cortical networks were numerically positive, with the CON and the cortical network average reaching significance at FDR p<.05. Conversely, the trajectory was in the opposite direction for subcortex (4/5 regions were negative), with VMM (comprising perirhinal regions whose functions are intimately linked to subcortex, see Methods) and the subcortical region average significantly decreasing over time. See Fig S5 for a simulated demonstration that the observed early negative conjunctive estimates indicate dominance of compositions over conjunctions, with increases in conjunctive estimates over time truly indicating progressive strengthening of those conjunctions. Panels **B-D** provide the compositional (single rule) results. Compositional codes were reliably present in cortex and subcortex, and emerged early. However, compositional codes were significantly stronger in cortex > subcortex (paired t-tests for all blocks, FDR p<.05). The dynamic trajectory analyses revealed a generally negative trend, with compositional codes weakening over time in both cortex and subcortex.

Further dissociation was revealed in the dynamic trajectories. Whereas conjunctive codes progressively strengthened in cortex (cortical average, trajectory t=2.70, p=.009), they progressively weakened in subcortex (subcortical average, trajectory t=-2.77, p=.008), and this difference was significant (paired t-test cortex > subcortex, t=3.52, p=.001). Contrastingly, compositional codes progressively weakened across cortex (mean across 3 rules, cortical network average, t=-4.22, p<.001) and subcortex (mean across 3 rules, subcortical region average, t=-3.78, p<.001). This profile is consistent with the hypothesized importance of strengthening conjunctive representations in cortical networks specifically over repeated task practice (Fig 1).

The Supplement (Fig S5) provides an expanded statistical explanation of the Fig 5 results, with a focus on correctly interpreting the early negative cortical conjunction estimates (Fig 5A) as arising from representational interference between the compositions. This references established statistical understanding of the interaction term in regression models, complemented by simulations with a known ground truth, to substantiate that the brain truly exhibits an increasing reliance on cortical conjunctions with practice.

The Supplement (Fig S6) also reports equivalent representational dynamics over learning when using an alternative, established measure of estimating compositional representations (cross-condition generalization performance; Bernardi et al., 2020). As detailed in the Supplement, we were precluded from applying an alternative method of estimating conjunctive representations (Kikumoto & Mayr, 2020) by inherent restrictions in our task design, specifically the lack of novel task repetitions (which would have invalidated the novelty of those tasks).

### Strengthening of cortical conjunctions is uniquely associated with behavioral improvement over practice

We hypothesized that the dynamic shift from compositional to conjunctive representations over task practice would be associated with behavioral improvement. We computed associations between the rule strength timecourses and block-to-block variation in reaction time (RT). The results in Fig 6 demonstrate that stronger cortical conjunctions were linked to behavioral improvement (quicker RT) with practice, particularly in the FPN (t=-4.11, p<.001). Conversely, associations between compositional strength and behavior over this extended timescale were generally in the harmful direction (slowing RT), for both cortex and subcortex, and especially for the sensory rule type. The findings are overall consistent with our hypothesis that cortical conjunctive strengthening is a key computational signature of behavioral optimization with practice.

**Fig 6.**
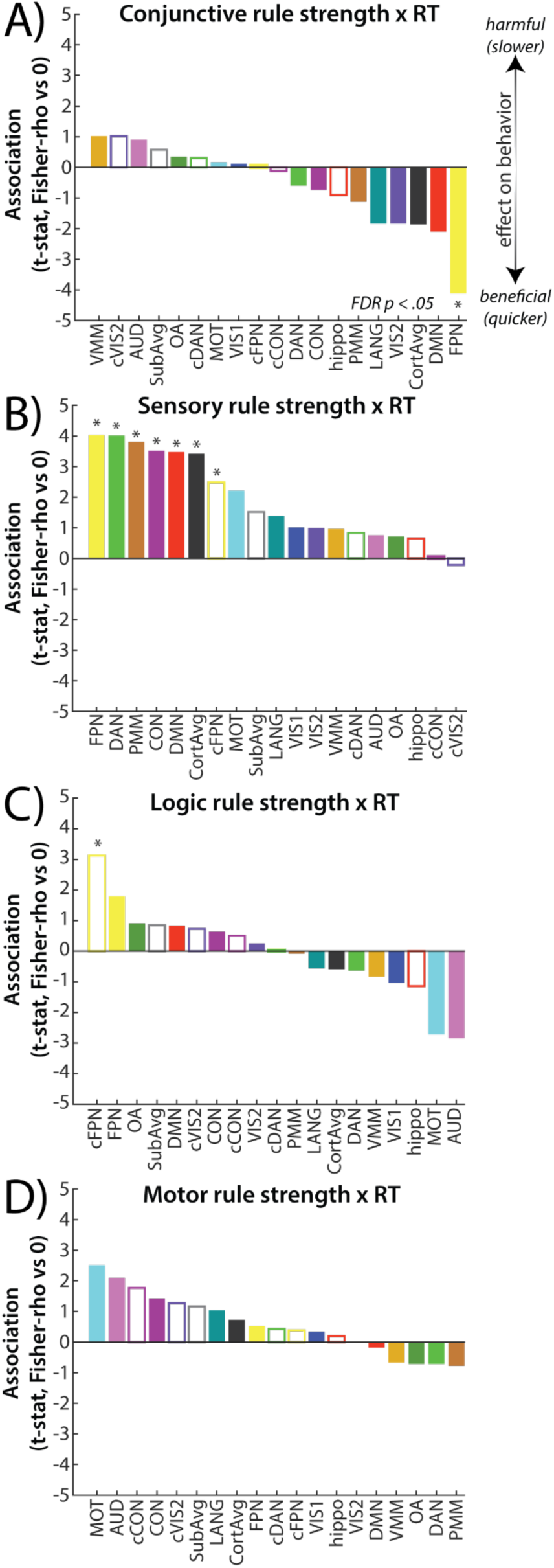
Strengthening of cortical conjunctions over learning is associated with behavioral improvement (quickening of RT). Panels display the Spearman correlation between rule representation strength and behavioral RT. Conjunctive strength was generally numerically associated with beneficial effects on behavior, reaching significance in the FPN cortical network (FDR p<.05). Conversely, maintaining stronger rule compositions over this timescale of extended practice generally yielded harmful behavioral associations (slowing of RT), with this effect most pronounced for sensory rules in the cognitive control networks (CCNs) and DMN.

This pattern of brain-behavior association held in the FPN even after regressing out linear temporal effects prior to computing the associations via partial correlation (FPN conjunctive-RT association: t=-2.16, p=.036; FPN sensory-RT association: t=2.99, p=.005). This was an exhaustive control for temporal confounds/artifacts, conducted in addition to removing linear trends and expanded noise regressors during preprocessing (see Methods).

Our use of methods of brain-behavior association that were sensitive to both linear and non-linear relationships (i.e. Spearman’s correlation) was validated by stepwise regression modeling of the behavioral RT timecourses (see Methods). This revealed significant behavioral prediction accuracy when entering the time predictor linearly (β=0.40; R^2^=0.21, p<.001 via Wilcoxon signed-rank versus 0) and non-linearly (power law; β=0.39; R^2^=0.15, p<.001). Contrasting R^2^ for linear > non-linear models via paired Wilcoxon: Z=3.44, p<.001. Critically, the combined model in which linear and non-linear predictors were entered simultaneously yielded numerically greater behavioral prediction accuracy (combined R^2^=0.21, p<.001) than the linear only model (paired Wilcoxon contrasting R^2^ for linear versus combined, Z=-1.84, p=.065), and significantly greater accuracy than the non-linear only model (Z=2.89, p=.004). These results suggest that both linear and non-linear temporal effects were present in behavior, which justified our use of Spearman’s correlation-based approaches in the brain-behavior association analyses.

### Strengthening of cortical conjunctions over practice is associated with stronger pattern separation

We hypothesized that one advantage of strengthening conjunctive representations with practice is to increase pattern separation, and consequently reduce cross-task interference. We therefore interrogated whether pattern separation (reduced neural task similarity over learning; Fig 3) was associated with the strength of conjunctive or compositional rule representations (Fig 5). For each region, we computed the Spearman’s correlation between the separate measures of neural task similarity and rule representation strength, estimated across Practice session blocks. The results in Fig 7 demonstrate that compositional weakening over this extended timescale was associated with improved pattern separation. This held across cortex/subcortex, and across the 3 single rule types (cross-rule cortex average, t=36.86; cross-rule subcortex average, t=14.16; both p<.001). Weakened subcortical conjunctions were also associated with stronger pattern separation (subcortex average, t=4.62, p<.001). Critically, conjunctions in cortex uniquely showed the opposite, beneficial association: stronger cortical conjunctions related to stronger pattern separation (cortex average, t= −19.14, p<.001).

**Fig 7.**
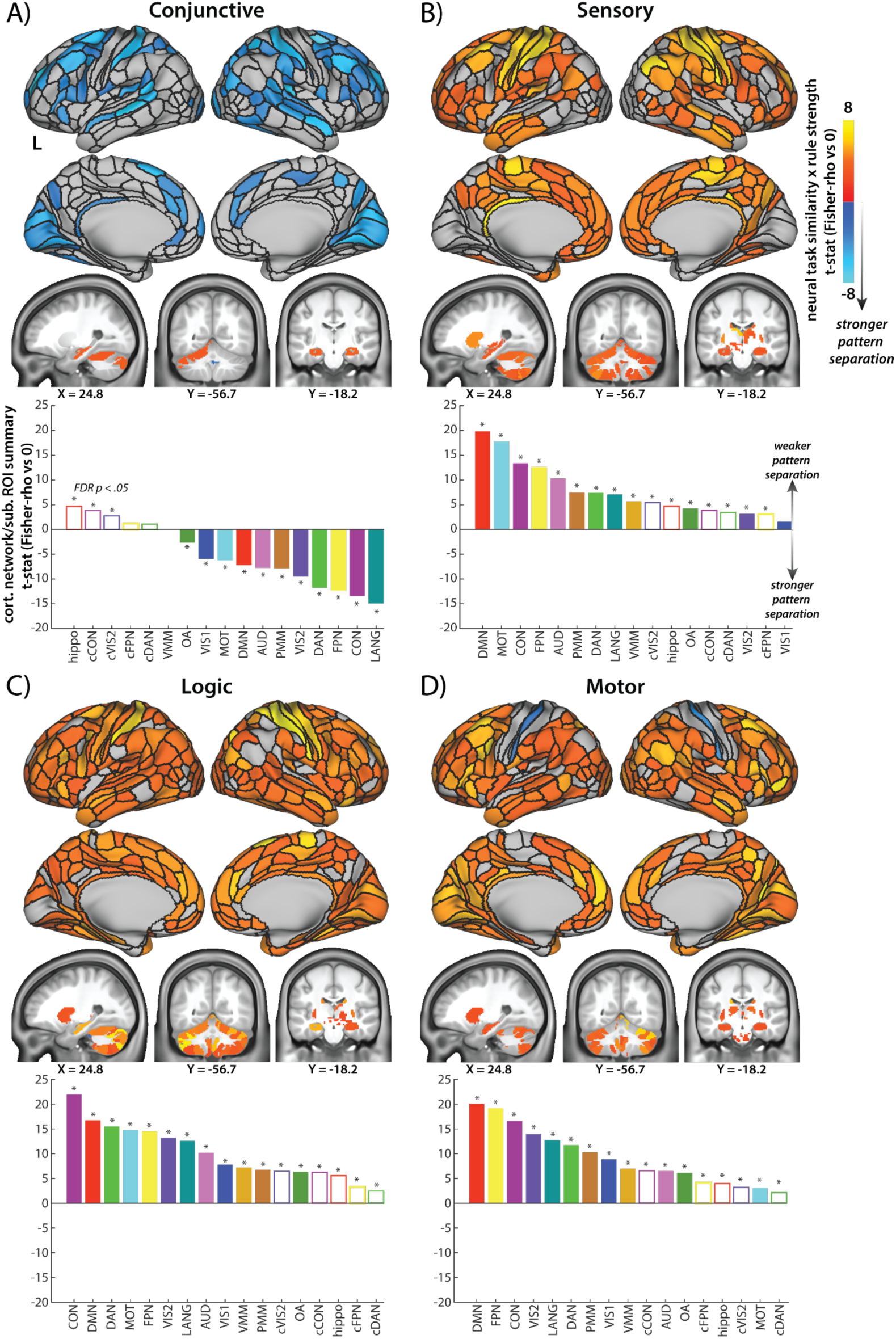
Cortical conjunctive strengthening is uniquely associated with stronger pattern separation over task practice. Panels depict Spearman correlations between neural task similarity (our index of pattern separation, Fig 3A) and conjunctive (panel **A**) or compositional rule strength (panels **B-D)**, thresholded at FDR p<.05. Panels plot regional results on the cortical surface (top row) and subcortical volume (middle row), as well as the average across cortical networks and subcortical ROIs (bottom row). Weakening of compositional rule strength over this timescale of extended practice was associated with decreasing neural task similarity (stronger pattern separation), and this association held across multiple regions in cortex and subcortex (including bilateral hippocampus, bilateral cerebellum, bilateral dorsal striatum, bilateral thalamus, right amygdala and bilateral brainstem). The direction of association between pattern separation and conjunctive strength varied for cortex versus subcortex. For subcortex, weakening conjunctive representations were related to decreasing neural task similarity (stronger pattern separation). For cortex, increasing conjunctive representations were beneficially related to decreasing neural task similarity (stronger pattern separation). Significance in the network/regional average plots across panels (denoted by asterisks) highlighted that the effects were spatially distributed, albeit with overall greatest prominence in the CCNs, DMN and higher sensory networks.

The overall spatial profile is again consistent with the hypothesized differentiation in function between cortex and subcortex over task learning. Critically, the evidenced beneficial relationship between cortical conjunctions and pattern separation was predicted by our theoretical framework (Fig 1), as a signature of effective practice that reduces representational interference.

### Cortical conjunctions continue to strengthen across a second learning session

If cortical conjunctions serve as a neural signature of the extent/robustness of task learning (per our hypotheses), then they should continue to strengthen with additional practice. Whereas preceding analyses focused on the first Practice session, here we estimated the strength of conjunctive representations for practiced tasks presented in the second Test session (separated from the first session by 1-7 days; see Methods).

Consistent with our predictions, the timecourses in Fig 8A revealed further strengthening of cortical conjunctions as practice progressed in the Test session. This visual impression was formalized via contrasts of rule strength over temporal epochs (see Methods). This revealed significantly stronger (more positive) cortical conjunctions for Test session epochs compared to Practice session epochs (early Prac < early Test, t=3.61, p<.001; early Prac < late Test, t=4.06, p<.001; late Prac < early Test, t=2.99, p=.005; late Prac < late Test, t=3.15,p=.003; all p values survived FDR p<.05 multiple comparison correction). The remaining contrasts were in the same direction of stronger cortical conjunctions with increasing practice, albeit non-significant (early Prac < late Prac, early Test < late Test, both p>.570).

**Fig 8.**
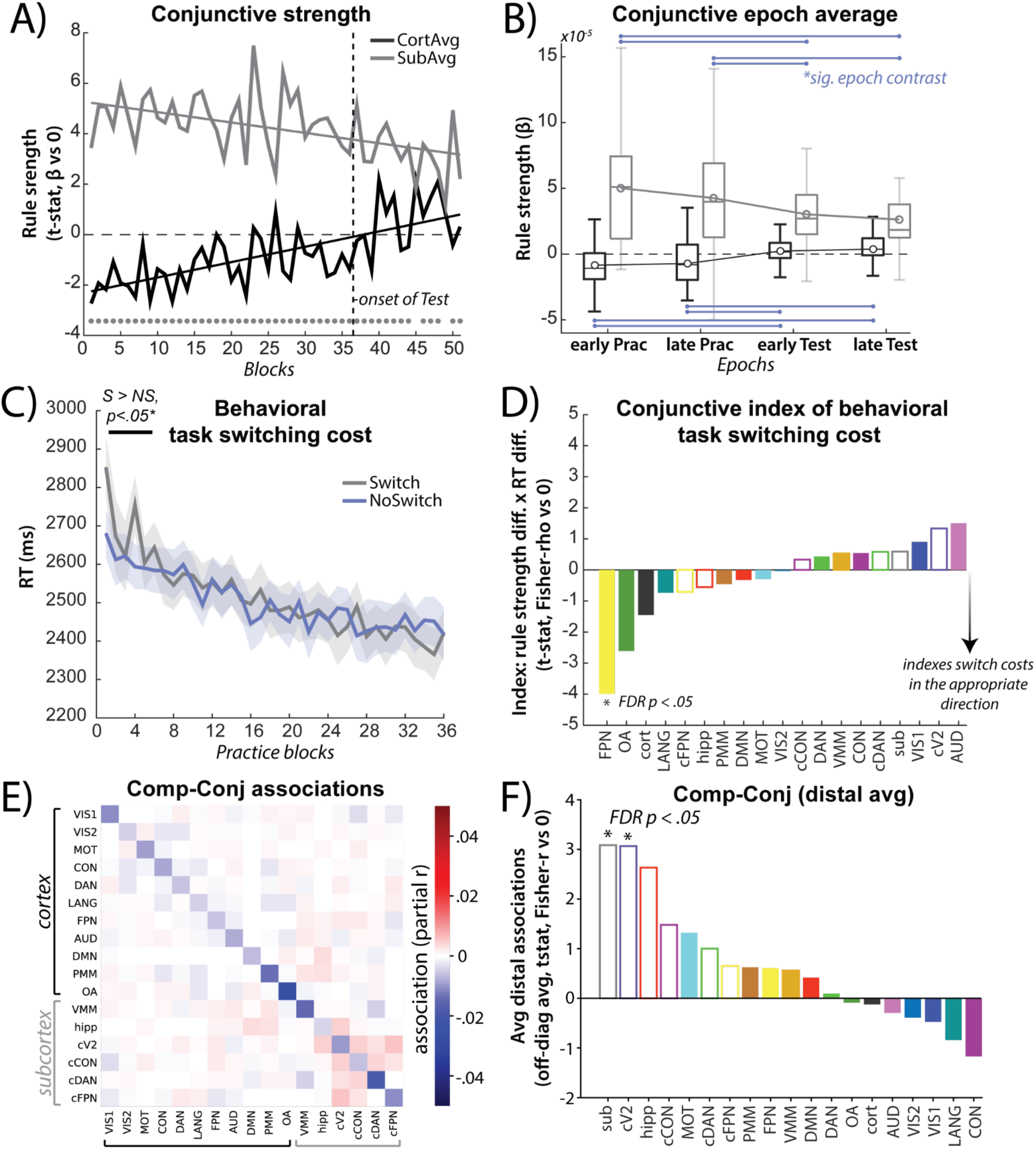
Cortical conjunctions strengthen into the second Test session, and index behavioral costs of task switching, but subcortex is critical for cross-region binding of rule information. A) Test session conjunction analysis: dynamics of conjunctive rule strength across the first Practice and second Test sessions (onset of the latter marked with a dotted line). Estimation of conjunctive strength and norms of plotting are the same as in Figure 5. Cortical conjunctions (black lines) continue to strengthen and subcortical conjunctions (gray lines) continue to weaken with practice. B) Test session conjunction analysis: epoch averages of the timecourse results in panel A (see Methods for details). Horizontal lines in each box represent the median, circles the mean, edges of each box the interquartile range, and whiskers the data range. Blue lines indicate significant paired t-test contrasts between epochs (FDR p<.05). These plots formalized the trends in panel A: cortical conjunctions were significantly stronger in Test versus Practice session epochs; subcortical conjunctions were significantly weaker in Test versus Practice epochs. C) Task switching analysis: behavioral cost of increased task switching. Reaction time was significantly slower for Switch compared to NoSwitch practiced tasks in early Practice session epochs. D) Task switching analysis: brain-behavior association. Differences in FPN conjunctive strength across conditions (NoSwitch > Switch) reliably indexed the behavioral cost of task switching (NoSwitch RT < Switch RT). E) Cross-region association analysis: group-averaged matrix, capturing associations of compositional and conjunctive rule strength dynamics between cortical networks and subcortical regions. The matrix reveals greater positive associations (red matrix elements) involving subcortical compared to cortical regions, reflecting greater subcortical sensitivity to binding of compositional rule information into conjunctions. F) Cross-region association analysis: averaging the off-diagonal elements in the panel E matrix, for each network/region, and contrasting against 0 via one-sample t-test. This graph summary confirmed that subcortical regions were most prominent in binding rule information.

Subcortical conjunctions in the Test session showed the opposite trajectory, progressively weakening as practice continued (Fig 8A). This was formalized in the epoch analysis (Fig 8B) as significantly weaker subcortical conjunctions in Test compared to Practice epochs (early Prac > early Test, t=3.11, p=.003; early Prac > late Test, t=3.64, p<.001; late Prac > early Test, t=2.27, p=.028; late Prac > late Test, t=2.62, p=.012; all FDR p<.05). The remaining contrasts were numerically in the same direction but non-significant (early Prac > late Prac, early Test > late Test, both p>.330). These findings reveal that the conjunctive dynamics observed in the first Practice session were extended with additional practice in the second Test session. The opposing trajectories between cortex and subcortex further supports our hypothesized division of labor between these systems in task learning. Critically, the results substantiate cortical conjunctive strength as a neural index of the extent of task practice, even as it extends across separated learning episodes.

For completeness, the Supplement presents compositional rule strength estimated in the Test session (Fig S7). This revealed transient compositional re-engagement at the start of the Test session, as a response to increased cognitive demand following the lapse in task performance, as reflected in behavior (Fig S8). The differing dynamic profile of compositions and cortical conjunctions is again consistent with our hypothesis that they underpin separate neural computations, albeit suggesting that compositional engagement occurs under general increase in demand, rather than exclusively under novelty.

### Behavioral cost of task switching is uniquely indexed by cortical conjunction strength

We leveraged another C-PRO2 manipulation to substantiate the link between cortical conjunction strength and the robustness of practiced task representations. Task switching demands were varied across the 4 practiced tasks: 2 tasks were presented under high switching (Switch condition), and the remaining 2 tasks under reduced switching (NoSwitch condition; see Methods). We predicted that the Switch condition would interfere with effective practice, leading to a behavioral cost relative to the NoSwitch condition, which would be indexed by the strength of cortical conjunctions.

We first verified that high task switching yielded the anticipated behavioral cost. Behavioral timecourses revealed slower RT for Switch compared to NoSwitch tasks early in the Practice session (epoch average over blocks 1-6; Fig 8C); Switch early epoch mean=2691.2 ms, NoSwitch early epoch mean = 2609.6 ms, paired t=2.40, p=.021. Accuracy also worsened for Switch tasks during this period (Fig S9A); Switch early epoch mean=66.85%, NoSwitch early epoch mean = 73.61%, paired t=2.24, p=.030. Late epoch Switch vs NoSwitch contrasts were non-significant (both p>.240). This profile is consistent with increased task switching disrupting effective practice.

We then assessed whether the strength of cortical conjunctions was sensitive to this switching cost. Conjunctive representations were estimated separately for Switch and NoSwitch conditions. The resulting timecourses (Fig S9B) revealed similar dynamic trajectories across Switch and NoSwitch, verifying that prior analyses that collapsed across these conditions didn’t occlude dramatic differences. However, the intercept visibly differed between switching conditions. We formalized these effects via a 2 (brain system: cortex, subcortex) x 2 (switch condition: Switch, NoSwitch) repeated measures ANOVA on the session-level (block-averaged) conjunctive representation strength. This yielded a significant main effect of system (Subcortex > Cortex, F(1,43) = 77.0, p<.001, η^2^g=0.413) and a non-significant main effect of condition (Switch > NoSwitch, F=2.67, p=.109, η^2^g=0.004). Critically, the system-by-switch condition interaction was significant (F=5.97, p=.019, η^2^g =0.009), reflecting a reliable reversal in the sign of the Switch vs NoSwitch effect between cortical and subcortical conjunctions. Planned paired t-tests revealed that cortical conjunctions were numerically stronger for NoSwitch tasks (Switch estimate (t) = −2.59, NoSwitch estimate (t) = −1.72; Switch > NoSwitch paired t = −0.77, p = .449), whereas subcortical conjunctions were significantly stronger for Switch tasks (Switch estimate (t) = 8.07, NoSwitch estimate (t) = 7.22, paired t = 2.48, p = .017). This dissociable profile again supports the division of labor between cortical and subcortical systems in task learning.

We then examined whether block-to-block variation in cortical conjunction strength was associated with block-to-block variation in behavioral switch costs. Figure 8D plots correlations between conjunctive rule strength differences (NoSwitch minus Switch) and behavioral RT differences (NoSwitch minus Switch) across Practice session blocks. The results in Fig 8D reveal that conjunctive strength in the FPN appropriately and uniquely indexed the behavioral costs of task switching (t= −3.99, p<.001; only network that survived at FDR p<.05). Follow-up analyses revealed FPN’s early conjunctive strength to be significantly greater for NoSwitch versus Switch tasks (NoSwitch estimate (t)=0.34, Switch estimate (t)=-2.09, paired t=2.14, p=.038), demonstrating sensitivity to the period of maximal behavioral cost. FPN’s brain-behavior association index also held when confined to this early epoch of maximal cost (Spearman’s correlation confined to blocks 1-6; t=-2.29, p=.027). This FPN index was evidenced even after regressing out the linear temporal effect before computing the index, to rule out potential temporal confounds/artifacts (partial t=-3.45, p=.001). This aligns with earlier brain-behavior associations in the FPN that were also robust to this control analysis.

For completeness, the supplement provides equivalent results for compositional rule strength (Fig S10), revealing overall modest differences across switching conditions, and no statistically reliable brain-behavior association index. This further substantiates the unique link between FPN conjunctions and practice-related behavioral improvement. The behavioral index captured more robust strengthening of conjunctions in the FPN under reduced task switching demands, which associated appropriately with behavioral improvement in that condition.

### Cross-region association of rule representation dynamics reveals subcortical binding of compositions into conjunctions

Preceding analyses interrogated rule representation dynamics separately within each region and rule representation type (compositions and conjunctions). In ensuing analyses, we examined profiles of cross-region and cross-rule associations, to reveal insight into network interactions underlying the task learning dynamics.

We used a regularized multivariate measure of cross-region association, inspired by recent advancements in functional connectivity estimation (Peterson et al., 2023; see Methods). This measure was computed for all pairwise combinations of rule strength timecourses (i.e. compositional-compositional, conjunctive-conjunctive, and compositional-conjunctive associations), across Practice session blocks. We hypothesized that compositional-compositional associations would primarily capture how the same kind of rule information was propagated across the brain, and that this would prominently involve cortical networks. Conversely, compositional-conjunctive associations would capture the transformation of different rule representation formats, and prominently involve subcortical regions. These predictions follow from the differential sensitivities of cortex/subcortex to compositions/conjunctions in prior analyses, and multiple memory systems theories linking subcortex to binding operations (Ranganath, 2010; Yonelinas et al., 2019). We did not have strong predictions regarding the cross-region conjunctive-conjunctive associations, but the highly specialized nature of these representations raised potential difficulties in registering them at the spatial scale of fMRI (versus e.g. the scale of neural populations).

The group-averaged matrix for compositional-compositional cross-region associations is provided in the Supplement (Fig S11A). This revealed generally positive compositional-compositional associations, which were more prominent in cortex than subcortex. The visual impressions were formalized by averaging the distal (off-diagonal) associations for each cortical network/subcortical region (Fig S11B), which revealed significantly greater associations for the cortical average compared to the subcortical average (paired t=16.6, p<.001). The conjunctive-conjunctive results (Fig S11C and S11D) revealed no significant cross-region association, suggesting that these were primarily local, and/or that distal associations were at too fine a scale to be registered with fMRI.

Fig 8E plots the critical compositional-conjunctive cross-region associations, which were visibly more positive for subcortex versus cortex. This impression was formalized by averaging the off-diagonal elements, which revealed significantly positive associations involving a cerebellar region (cV2) and the subcortical average (at FDR p<.05; hippocampus was significant at uncorrected p<.05). The subcortical average was also significantly greater than the cortical average (paired t=3.48, p=.001). The findings substantiate our prediction that subcortical regions would be prominently involved in binding task-general compositions into more specialized conjunctions. The pattern of results again highlights a division of labor between cortex (propagating the same kind of compositional information across the brain) and subcortex (binding compositions into conjunctions).

## Discussion

We assimilated concepts from separate neuroscientific subfields of long-term memory, language, context-dependent decision-making and cognitive control to test a neurocomputational framework of task learning (Fig 1). This formalized a dynamic shift from compositional to conjunctive representations over practice. We developed methods to quantify this shift in the functioning human brain as multi-task learning progressed (Fig 2A). The findings support our hypotheses: compositional representations are engaged early in learning to facilitate transfer of prior knowledge to novel contexts, while progressive strengthening of cortical conjunctions drives effective practice (behavioral optimization and reduced interference). Shifting from task-general compositional representations to task-specific conjunctive representations hence serves as a key neural underpinning of successful practice.

We firstly demonstrated decreases in neural representational similarity with task practice, extending pattern separation computations beyond the spatial memory and visual object recognition domains of typical study (Bakker et al., 2008; Brunec et al., 2020; Marr, 1971) to cognitive task learning. We then evidenced a precise computational basis for such changes: increasing conjunction (binding) of each task’s rule components. Conjunctions have long been considered an important neurocomputation of long-term memory (Eichenbaum, 2013; Marr, 1971; Radulescu et al., 2021; Shimamura & Wickens, 2009; Sutherland & Rudy, 1989; Wickelgren, 1979), and are central to complementary learning systems theory (O’Reilly & Rudy, 2001). Here we used machine learning to dynamically quantify the strength of conjunctive and compositional codes in empirical data, taking inspiration from recent analytic innovations in context-dependent decision-making (Bernardi et al., 2020; Dang et al., 2021; Kikumoto & Mayr, 2020; Rigotti et al., 2013). Whilst this literature has demonstrated the importance of conjunctions in within-trial dynamics underlying behavior, here we highlight their role in optimizing human behavior at the longer timescale (minutes/hours) of task learning.

Conversely, compositional representations, previously highlighted as a basis for flexible knowledge transfer (Bernardi et al., 2020; Cole et al., 2011; Cole, Laurent, et al., 2013; Collins & Frank, 2013; Lake et al., 2015; McClelland & Rumelhart, 1985; Yang et al., 2019), were prominent early in learning when tasks were novel, and weakened thereafter. The supplement (Fig S7 and S8) also provides evidence for a more general link between compositional representations and high demand conditions (novelty and lapses in task performance). Our findings add to emerging evidence across subfields of a fundamental trade-off between encoding and retrieving generalities of experience (compositional coding), and encoding and retrieving specifics of experience (conjunctive coding; (Behrens et al., 2018; O’Reilly & Rudy, 2001; Radulescu et al., 2021; Reagh & Ranganath, 2023; Whittington et al., 2022). This trade-off provides a neurocomputational underpinning to the transition from controlled to automatic processing (Chein & Schneider, 2005; Cole et al., 2010; Mohr et al., 2016; Schneider & Chein, 2003).

We consistently observed differential involvement of cortex and subcortex in the interplay between compositional and conjunctive codes. In the present timeframe (minutes/hours, relatively early in learning), cortex more strongly represented compositions, whereas subcortex more strongly represented conjunctions. This substantiates the division of labor posited by complementary learning systems theory (McClelland et al., 1995; O’Reilly et al., 2014; O’Reilly & Rudy, 2001), whilst adding computational detail through distinguishing conjunctive and compositional codes (the latter being neglected in the original theory). Elucidating representational dynamics was key in specifying this division of labor: compositions were engaged early in both cortex and subcortex and then weakened; conjunctions were engaged early in subcortex and then marginally weakened, whilst conjunctions in cortex slowly strengthened. The slow strengthening of cortical conjunctions was uniquely associated with behavioral improvement and pattern separation, highlighting it as a key signature of effective task practice. Importantly, this strengthening continued into a second task learning episode (Fig 8A-B), and was distinct from dynamics of compositional representations in this period (Fig S7). This provides empirical support for the computational tenets of multiple memory systems theories, which were extended here to characterize learning in the higher-order task rule domain.

Our findings also linked rule representation dynamics to behavioral costs of task switching (Flesch et al., 2018; Musslick & Cohen, 2021; Sakai, 2008; Xu et al., 2024; Yeung et al., 2006). Increased task switching during practice worsened behavior (Fig 8C, Fig S9A), and these behavioral costs were appropriately and uniquely indexed by FPN conjunctive strength (Fig 8D, Fig S10). Our findings suggest that task switching disrupts the time-intensive (and behaviorally critical) process of strengthening cortical conjunctions over practice, leading to increased representational interference and behavioral costs. This contrasts with learning under reduced task switching conditions, which allows more uninterrupted time for cortical conjunctive strengthening. These findings again portray cortical conjunctive strengthening as a key neurocomputational signature of effective learning, with relevance in explaining performance limitations due to task switching.

Beyond the clear differentiation between cortex and subcortex over learning, we also observed differentiable large-scale network involvement. Network-averaged results consistently highlighted regions within the DMN, CCNs and higher-sensory networks as showing the representational dynamics most prominently. Whilst the memory function of the DMN is increasingly well-established (Kaefer et al., 2022), a role for CCNs and higher sensory networks in learning computations (e.g. pattern separation) has recently been acknowledged (Amer & Davachi, 2023), building on earlier work linking these networks to general cognitive control functions (Cocuzza et al., 2020; Cole, Reynolds, et al., 2013; Mill et al., 2022), and control operations impacting long-term memory (Dobbins et al., 2002; Mill et al., 2015, 2016; Simons & Spiers, 2003). Future work will be necessary to elucidate the precise nature of interactions between the DMN, CCNs and sensory networks that give rise to the representational dynamics we observed (de-Wit et al., 2016; Ito et al., 2019). The centrality of CCNs to these interactions is supported by FPN conjunctions showing the strongest associations with behavior, and behavioral costs of task switching.

We also examined cross-region synchronization of representational dynamics, finding cortical networks to be prominent in propagating compositional rule information (Fig S11), whereas subcortical regions were prominent in transforming/binding compositions into conjunctions (Fig 8E-F). These results substantiate the profile of hypothesized cortical-subcortical interaction (Fig 1A). The Supplement (Fig 56) presents simulations consistent with a network mechanism wherein early task-general compositional engagement is progressively sparsified around task-relevant sub-circuits, thereby boosting efficiency of information processing from stimulus to response over practice.

Future work will be required to target network interactions underpinning the observed representational dynamics more formally, which may draw on recent empirical neural network modeling approaches (Cole et al., 2016; Hearne et al., 2021; Ito, Yang, et al., 2022; Mill et al., 2022). Such methods permit adjudication between candidate network models of task learning in empirical data, which could illuminate the mechanism of interaction between cortex and subcortex, and the degree to which they operate independently or synergistically (O’Reilly et al., 2014; Schapiro et al., 2017; Squire et al., 2015; Whittington et al., 2022).

Whilst we believe our approach to fMRI preprocessing and expanded nuisance regression was rigorous enough to rule out major artifacts (see Methods), future research seeking convergence of our findings across different neural modalities (e.g. source EEG/MEG) would capitalize on the unique artifacts in these data to increase confidence. It will also be important to study representational dynamics beyond the limited window of task learning studied here (minutes/hours of practice), and towards a more extended timescale (hours/days of practice). This is especially as we may hone our efficiency in performing critical tasks over a lengthy timescale (days/months/years) in daily life.

Whereas the current limited timescale was appropriate in capturing the retrieval of compositions (promoting minimal competency in specific tasks) and encoding of new cortical conjunctions (promoting efficiency in specific tasks), a more extended timescale may reveal how new compositions are encoded (providing new building blocks for task-general transfer; Sun et al., 2023).

Although the present work provides important empirical substantiation of the long-theorized division of labor between cortex and subcortex, future work seeking to separate roles amongst subcortical regions would help advance beyond the relatively coarse “cortex versus subcortex” distinction. Whilst the involvement of hippocampus in learning has been widely scrutinized, cerebellum has received less interest, despite being linked to relevant functions of procedural motor skill learning (Beukema & Verstynen, 2018; Ferrari et al., 2018) and behavioral automaticity (Lang & Bastian, 2002; Nixon & Passingham, 2000). Our results suggest a precise computational basis for cerebellar contributions to learning, via early encoding of rule compositions and conjunctions. These computations are supported by cerebellum’s dense bidirectional connectivity with MTL and cortical regions (Buckner, 2013; Z. Gao et al., 2018; Strick et al., 2009). Future work that manipulates reward and expectation will be useful in probing differentiation between MTL and cerebellum, which may also generally benefit from the improved spatial resolution of high field fMRI.

To conclude, we elucidated specific cortical and subcortical computations underlying changes to neural representational geometry over task learning. We believe the neurocomputational signature identified here – progressive cortical conjunctive strengthening that drives pattern separation and behavioral improvement – is a general learning mechanism, with relevance to multiple neuroscientific subfields. Future examination of the network underpinnings and generality of this mechanism has potential to advance fundamental understanding of learning in multiple contexts.

## Methods

### Subjects

Usable fMRI data were collected from 44 subjects (age mean=22.25 years, range=18-36; 23 self-reported female), out of a total of 46 (one subject was excluded due to a computer malfunction, and another for failing to use the response pad correctly). The sample was recruited from Rutgers University-Newark and neighboring communities, with the inclusion criteria of right-handedness, absence of current or past psychiatric or neurological illness, and absence of standard MRI contraindications. Prior to MRI data collection, a separate sample of 34 subjects (age mean=25.74 years, range 19-40; 18 self-reported female) was included in a behavioral study that validated the stimuli created for the task (see next section for details). All subjects gave informed written consent prior to participating, following ethical guidelines from the Rutgers University Institutional Review Board (IRB).

### C-PRO2 stimulus creation and task design

Our aim of elucidating general neural principles of cognitive task learning informed substantial updating of the permuted rule operations (PRO) paradigm. Whereas previous versions used abstract word stimuli (Cole et al., 2010) or concrete multi-sensory stimuli (C-PRO; visual lines and auditory tones; Ito et al., 2017), the present concrete version (C-PRO2) used more complex, naturalistic and variable multi-sensory stimuli. The sensory dimensions of these stimuli (described below) were chosen based on prior literature suggesting they would elicit strong and spatially distinct neural representations (Epstein & Kanwisher, 1998; Formisano et al., 2008; Kanwisher et al., 1997; Lafer-Sousa et al., 2016).

Visual stimuli were sampled from established image databases, and varied on the following dimensions: young versus old faces (CAL/PAL face database; Minear & Park, 2004), and indoor versus outdoor scenes (SUN database; Xiao et al., 2010). The young (18-32 years of age) and old (64+ years) faces were taken from the Ebner (Ebner, 2008) update of the original CAL/PAL database, which sub-sampled to emotionally neutral expressions and color images, and additionally processed the images to remove eye-catching non-facial features (e.g. jewelry) and standardized low-level visual features (luminance and color, across background and clothing). To further reduce low-level feature variability, we used Adobe Photoshop to standardize image size and opacity (50%), for both faces and scenes. Faces were also cropped to remove hair, which could serve as an unwanted low-level visual cue for the young versus old face category judgment. The final composite visual stimuli consisted of faces overlaid centrally on scenes (Fig 2A), capitalizing on the reduced opacity of the constituent images.

Auditory stimuli were sampled from the ACCENT database (Weinberger, 2016), which contains audio recordings of the same 30-second short story read by multiple human speakers of varied gender, nationalities and accents. Our sampling was confined to US American accents to match the location of the study, with an equal proportion of normatively masculine (male) and normatively feminine (female) voices. The continuous speech recordings were then manually segmented into non-overlapping 3-word phrases using the PRAAT software (Boersma & Weenik, 2023). Further processing to control for low-level auditory feature variability included adding a 50 ms fade in/out period at the start/end of each sound (to correct for acoustic clicks), equating the duration of all sound files to 800 ms (without affecting pitch, using PRAAT’s PSOLA algorithm) and equating the intensity of all files to 75 dB. Reversed versions of each segmented audio file were also created to serve as the alternate response category. Hence, the final set of auditory stimuli varied across two dimensions: male versus female speaker, and word (lexical) vs non-word (non-lexical i.e. reversed) content.

To ensure that stimuli were reliably discriminable across all 4 sensory dimensions, a behavioral validation study was conducted on a preliminary set of 100 faces, 100 scenes, 100 word phrases and 100 non-word phrases (each with an equal proportion of relevant sub-categories e.g. 50 young and 50 old for the face condition). This study presented composite visual or composite auditory stimuli on individual trials, with subjects cued to categorize one dimension (e.g. “is the face in the image old or young?”), with the secondary dimension (in this case: scenes) randomly selected. Each stimulus was presented twice per subject, with categorization accuracy averaged across presentations and subjects. These data were first used to eliminate non-discriminable stimuli (any item yielding accuracy below 50% chance), and extreme high/low accuracy items were then iteratively eliminated from each of the 4 tasks to reduce accuracy differences between categories for each dimension. This procedure resulted in 78 stimuli for each of the 4 dimensions, which were reliably discriminable relative to chance (average accuracies across all stimulus categories were > 87%), and did not differ between categories (e.g. male voices were not significantly more discriminable than female voices; all between-category 2-sample t-tests non-significant at p > 0.16). Stimuli for the main study were created by permuting across these 78^4 possible stimulus compositions (pseudo-randomly at the group level to maximize stimulus variety), thereby constituting a highly complex and variable stimulus set that balances the greater naturalism of recent movie-watching studies (Sonkusare et al., 2019) with greater experimental control of simpler tasks. The goal of increasing stimulus variability was to increase the likelihood of results from this paradigm generalizing to a wide variety of contexts. These stimuli were included as the updated sensory dimension in the full C-PRO2 task (Fig 2A).

To provide an overview of the design, C-PRO2 presents 64 unique tasks per subject, based on permuting 3 rule types (logic, sensory and motor). Each rule type has 4 levels each, meaning that there are 12 rules total (see Fig 2A for the list of rules). Hence, each of the 64 tasks is defined by its unique combination of 3 rules, one each sampled from the rule types (one sensory, one logic and one motor rule per task). For each subject, 4 tasks were repeatedly presented (practiced task condition), with the remaining 60 tasks presented just once (novel task condition). Each of the 12 rules were featured once across the 4 practiced tasks assigned to a given subject, to ensure that individual rules were equally practiced overall. Practiced tasks were counterbalanced across subjects, such that all 64 tasks were included as practiced tasks across the group. There were 16 practiced task combinations total (16 * 4 = 64), meaning practiced tasks were reused every 16 subjects. Pre-scan task instructions were kept minimal to preserve learning effects to be tracked in the scanner.

The format of an example task block is depicted in Fig 2A. Each block began with instruction cue screens that conveyed the rules for the ensuing task, followed by three trials in which subjects applied those rules. Each trial presented two multi-sensory stimulus probes, comprising visual (young/old faces superimposed on indoor/outdoor scenes) and auditory (vocalizations varying in lexical/non-lexical content and normative female/male speakers) stimuli, presented concurrently. Subjects attended to the relevant stimulus dimension based on the cued sensory rule, formed their decision based on the cued logic rule, and responded based on the cued motor rule. How subjects needed to integrate across the various rules to perform an example task trial is detailed in the text above the Probe 1 and Probe 2 events in Fig 2A. To clarify the response format, for a task with the motor rule “LEFT INDEX”, subjects would make a “TRUE” decision by pressing their left index finger, and a “FALSE” decision by pressing their left middle finger. That is, subjects were instructed beforehand to make TRUE/FALSE decisions for a given task by using opposing fingers of the same hand (as cued by the motor rule for that task).

Feedback on performance accuracy was provided at the end of each 12-block run. The presentation order of the task rule instruction screens was counterbalanced across subjects, with a consistent order maintained for a given subject across Practice/Test sessions. Permuting this general task format across practiced/novel conditions, and across the logic/sensory/motor rule dimensions, thereby constituted a flexible multi-task paradigm. This firstly provided the necessary compositional task rule structure to test our hypotheses (probing the tradeoff between compositional and conjunctive codes over learning), and also allowed for more generalizable inferences on the neural basis of task learning than standard single-task paradigms.

### Experimental procedure

Subjects participated in three sessions, each separated by 1-7 days: fMRI Practice (task fMRI during repeated presentation of the 4 practiced C-PRO2 tasks, intermixed blockwise), fMRI Test (task fMRI during presentation of the 4 practiced tasks and the 60 novel tasks, intermixed blockwise), and EEG Test (EEG acquired during an identical repeat of the fMRI Test session). The third session involved dense-array EEG data recordings (using the EGI HydroCel Geodesic Sensory Net; 1000 Hz sampling rate, 256 sensors, Cz reference electrode), which were not analyzed in the present report. 15 minutes of resting-state data were also acquired in each session (pre-task for the fMRI sessions and post-task for the EEG session). The fMRI Practice session involved 12 runs of 12 blocks, amounting to 144 blocks total (4 practiced tasks, each repeated for 36 blocks). The fMRI Test session involved 10 runs of 12 blocks, amounting to 120 blocks total (each practiced task repeated for 15*4 = 60 practiced blocks, and each novel task presented once = 60 novel blocks). Analyses in the present report primarily focused on the fMRI Practice session. The fMRI Test session was used here to derive templates for the rule representation strength analysis (see later Methods section). More extensive analysis of the fMRI Test and EEG Test sessions is planned for future studies. All task sessions were programmed and presented using E-Prime 2.

Note that in the Practice session, the order of presentation of the 4 practiced tasks was controlled, with 2 tasks assigned to a “Switch” condition (∼50% likelihood of switching between them within a 12-block run) and 2 tasks assigned to a “NoSwitch” condition (no switching within a 12-block run). To simplify the majority of our analyses, we collapsed across switch/no switch conditions when analyzing the practiced tasks. We also conducted analyses focused on this task switching manipulation (see later Methods section, Fig 8C-D, Fig S9, Fig S10). For the Test session, the first 12-block run always presented the 4 practiced tasks (with the intention of reactivating the learned representations from the Practice session), with novel tasks randomly intermixed with the practiced tasks in subsequent runs, and the last run always presenting novel tasks only (to ensure all were presented despite the surplus of novel compared to practiced tasks).

### fMRI acquisition

Data were acquired by the 3T Siemens Trio MRI scanner at the Rutgers University Brain Imaging Center (RUBIC). Whilst all subjects underwent whole-brain multiband echo-planar imaging (EPI) sequences with a 32-channel head coil, enforced replacement of scanner hardware (specifically, the head coil) that occurred during data collection forced us to change the fMRI sequence parameters. Subjects 1-18 had the following sequence: TR = 785 ms, TE = 34.8 ms, flip angle = 55°, Bandwidth 1924/Hz/Px, in-plane FoV read = 208 mm, 72 slices in axial orientation, 2 mm isotropic voxels, multiband factor = 8. Subjects 19-20 had the following sequence: TR = 785 ms, TE = 37.2 ms, flip angle = 55°, Bandwidth 1924/Hz/Px, in-plane FoV read = 208 mm, 56 slices in axial orientation, 2 mm isotropic voxels with a 1mm slice gap, multiband factor = 8. Subjects 21-46 had the following sequence: TR = 785 ms, TE = 37.2 ms, flip angle = 55°, Bandwidth 1924/Hz/Px, in-plane FoV read = 224 mm, 64 slices in axial orientation, 2.2 mm isotropic voxels, multiband factor = 8. The same fMRI sequence was maintained across all rest/task acquisitions and across Practice/Test sessions for a given subject. Throughout, the same anatomical T1 and T2 MRI scans were collected during the Test session (both with 0.8 mm isotropic voxels). Spin echo fieldmaps were acquired at the beginning of Practice and Test sessions to enable correction of distortions in the functional and structural images. The acquisition order for the Practice session was: fieldmaps, resting-state (15 mins), 12 runs of C-PRO2 (practiced tasks only, ∼7 mins each). The order for the Test session was: fieldmaps, T1, T2, resting-state (15 mins), 10 runs of C-PRO2 (practiced and novel tasks, ∼7 mins each).

### fMRI preprocessing

fMRI data were preprocessed using the Human Connectome Project (HCP) minimal preprocessing pipeline, which has been described in detail previously (Elam et al., 2021; Glasser et al., 2013). Briefly, the pipeline performs reconstruction and segmentation of the anatomical images, followed by functional image distortion correction (using the acquired fieldmaps), motion realignment, volume-to-surface mapping, spatial normalization to a standard MNI surface template and cross-subject surface alignment using multimodal structural/functional MRI information (MSM-All; Glasser et al., 2016).

The HCP pipeline outputs preprocessed CIFTI grayordinate fMRI timeseries (for ∼60k 2-D cortical surface vertices, and ∼30k 3-D subcortical voxels) in standard MNI space, which were then submitted to a general linear model (GLM). Separate GLMs were estimated for the fMRI Practice and fMRI Test sessions. We adapted recent best practice recommendations for fMRI nuisance regression of resting-state data (Ciric et al., 2017) to optimally remove artifacts from our task data. Referencing the resting-state literature was motivated by prior reports highlighting greater propensity for artifact contamination in these data (versus time-locking to manipulated task events; Huijbers et al., 2017; Laumann et al., 2016), leading to development of more rigorous artifact controls than standard task fMRI approaches. Nuisance regressors included in the task GLM were the 6 standard motion parameters, and physiological regressors specified per the aCompCor procedure (first 5 principal components estimated within white matter and ventricular masks; Behzadi et al., 2007). By sensitively estimating signals from predominantly non-neuronal areas, the aCompCor regressors have been shown to be effective in reducing a host of physiological (e.g. cardiac, respiratory) and technical confounds in fMRI, when compared to regressing out the averaged white matter/ventricular signals, or regressing out externally recorded physiological signals (Behzadi et al., 2007; Caballero-Gaudes & Reynolds, 2017). Derivatives and quadratics for the 6 motion and 10 physiological regressors were also included, meaning that a total of 64 nuisance regressors were modeled. This constitutes a more rigorous and expanded approach to task fMRI nuisance regression than is typically conducted, which aimed to maximally reduce physiological, technical and motion artifacts in our analyses. Critically, each acquired fMRI run was linearly detrended before being entered into the GLM, which removed trend/drift artifacts. We also conducted an exhaustive partial correlation control analysis to rule out any residual temporal confounds in our brain-behavior association results (see later Methods section).

For both Practice and Test sessions, separate task regressors for each presented block were modeled, as boxcars convolved with the SPM canonical hemodynamic response function, spanning the onset of the first instruction screen to the offset of the last trial for that block. Our decision to model fMRI block activations that collapsed across instruction and trial phases was driven by a desire to capture rule representation processes that were evoked across both these phases. For each vertex/voxel, this yielded 144 task beta activation estimates spanning presentation of the practiced tasks across the Practice session, and 120 task beta activation estimates for presentation of the practiced and novel tasks in the Test session.

### Parcellation into functional regions and networks

Generally, the units of our fMRI analyses were task activations estimated for cortical vertices and subcortical voxels (both at 2mm scale), spanning the whole brain. We used within-region vertex/voxel activations as features in our multivariate analyses. Region boundaries were identified from established functional atlases: 360 cortical regions of interest from the Glasser surface atlas (Glasser et al., 2016), and 358 subcortical regions from the CAB-NP atlas (Ji et al., 2019). The use of these established regional atlases allowed for a principled means to define functionally relevant regions, with greater generalizability given that these definitions were established in previous held-out data. This also fulfilled our desire to include parcellations of both cortex and subcortex in our analyses, which addressed general trends in memory and decision-making subfields of excluding cortex and subcortex respectively. The CAB-NP also provides 12 functional network affiliations for all 718 regions (see Fig 3C), which were used to summarize the regional effects at the network level and aid functional interpretations. The expanded names of network labels provided in figures are as follows: AUD (primary auditory), CON (cingulo-opercular network), DAN (dorsal attention network), DMN (default mode network), FPN (fronto-parietal network), LANG (language i.e. higher auditory network), MOT (motor), OA (orbito-affective), PMM (posterior multi-modal), VIS1 (primary visual), VIS2 (higher visual), VMM (ventral multi-modal).

### Estimation of neural task similarity (pattern separation measure)

To test our pattern separation hypotheses, we used an approach inspired by representational similarity analysis (RSA; Diedrichsen & Kriegeskorte, 2017; Haynes, 2015) which we term neural task similarity. The approach is schematized in Fig 3A. For a given region, the Spearman rank correlation was computed between the vectorized activations of its within-region features (vertices for cortex, voxels for subcortex), across all 4 practiced tasks. This yielded 6 pairwise rho values for each block, which were averaged to provide a summary similarity value for that block. Repeating over blocks populated the neural task similarity timecourse for that region. Similarity timecourses were estimated for practiced tasks in the first (Practice) and second (Test) sessions. The Practice session timecourses were used to quantify dynamic trajectories over time (Fig 3B and 3C), as well as to examine associations with task behavior (Fig S3) and compositional/conjunctive rule strength (Fig 7, detailed below).

The Test session timecourses were averaged over blocks, and contrasted between practiced and novel tasks in the Test session (Fig 3B). For the Test session practiced tasks, this involved averaging neural task similarity over all pairwise practice-to-practice task similarity values, over all 15 block presentations. For the Test session novel tasks (which were only presented once), this involved averaging over all pairwise novel-to-novel task similarity values, after excluding any novel-to-novel task pairs that shared a rule. The latter step was important to ensure that similarity was not spuriously inflated by rule overlap between the novel tasks, thereby maintaining parity with the practiced tasks that by design did not share rules. For example, the similarity estimate between the novel task “YOUNG FACE-SAME-LEFT INDEX” and another novel task “YOUNG FACE-BOTH-LEFT MIDDLE” was excluded from the overall novel similarity average, due to sharing the “YOUNG FACE” sensory rule. The resulting averaged practiced task similarity and averaged novel task similarity for each region was then contrasted via paired t-test. Note that the number of data points for practiced and novel tasks entering this analysis was balanced, given that both conditions were presented 60 times each at Test.

For the results in Fig 3B, an analytic conjunction approach was used, in which the practiced < novel task similarity contrast (i.e. regions showing reliably stronger pattern separation for practiced versus novel tasks) was used to filter the Practice session trajectory analysis (i.e. any region showing a reliable positive or negative neural similarity trend over time). This analytic conjunction approach helped isolate regions that showed the strongest pattern separation effects of task learning, whilst further ruling out any contamination by temporal artifacts.

### Quantification of compositional versus conjunctive rule strength

To interrogate precise changes in neural representational geometry that occurred during task learning, we developed a machine learning approach to quantify the relative contributions of compositional (single rule) versus conjunctive rule representations over time (Fig 4A). This took inspiration from other multivariate approaches developed in context-dependent decision-making field (Bernardi et al., 2020; Kikumoto & Mayr, 2020). For a to-be-analyzed practice task (in the Practice session), we created compositional representation templates for each constituent rule (one sampled each from sensory, logic and motor rule types) by averaging over relevant blockwise activations in the held-out Test session. Note that availability of the novel tasks in the Test session was critical in enabling estimation of the compositional rule templates. For example, estimating the compositional representation for the “YOUNG FACE” rule (Fig 2A) with only access to the practiced tasks would result in a template that confounded the rule representation with the task representation, given that this rule was only presented in one practiced task context in the first Practice session. With access to the novel tasks in the second Test session, which presented “YOUNG FACE” in multiple distinct novel tasks, a compositional template for this rule could be created by averaging over all presentations of that rule in the novel tasks. This allowed estimation of compositional rule templates that were not confounded by the practiced task representations.

Finally, a conjunctive template was created by multiplying the 3 compositional rule templates together, akin to a non-linear interaction term. We then fit a multiple linear regression model with all 4 template terms (spatial regressors) as predictors to the regional task activation vector for a given block:

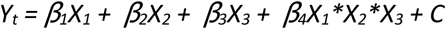

where *Y_t_* is the activation vector for the practiced task on that block, X1-3 are the single rule templates for sensory, logic and motor rules, and X_1_*X_2_*X_3_ is the rule template for the conjunctive rule (non-linear interaction term). Coefficients ꞵ_1-4_ therefore quantify the strength of each rule type in the activity representation evoked in that block. Note that we excluded practiced tasks presented in the Test session from data that were averaged to create each rule template, so as to prevent any unintended biases (e.g. inflating detection of the conjunctions, which likely were more strongly evoked in those practiced tasks).

We used robust regression to estimate the beta coefficients, given its improved resilience against outliers compared to regular least squares regression (Holland & Welsch, 1977). This was implemented via Matlab’s “robustfit” function, with default parameters (bisquare weight function, tuning constant=4.69). The resulting beta coefficients captured the unique strength of each rule representation during each block. Repeating over blocks and practiced tasks yielded timecourses of rule representation strength for each subject. The timecourses were averaged across the 4 practiced tasks, with the process repeated for each region to derive subject-level rule strength estimates.

Note that by estimating rule representation templates in held-out data, which are then fit to first-level regional activation patterns, our approach could be conceptualized as estimating/fitting spatial encoding models (Diedrichsen & Kriegeskorte, 2017; Ito et al., 2019). Encoding models are often considered to provide stronger tests of representational hypotheses than decoding models or representational similarity analysis, given the specification of particular spatial activation patterns as hypotheses rather than more abstract decoding hyperplanes or more vague representational similarity trends. Another beneficial feature of our approach is that we estimated the representation templates using fMRI data from an entirely different session, thereby going beyond previous calls for “cross-run” multivariate analyses, towards implementing a “cross-session” approach. This addresses temporal proximity confounds that have been previously highlighted in representational geometry analyses (Alink et al., 2015; Cai et al., 2019).

For the overall rule strength results in Fig 4B, we averaged the subject-level rule estimates across time (blocks), before conducting group-level random effects statistics. These results informed selection of subcortical regions-of-interest (ROIs) for ensuing analyses that preserved the block-to-block dynamics (Fig 5-8). Selected ROIs were subcortical regions showing the strongest pattern separation and conjunctive strength effects, with regional significance > 0 in both the neural task similarity (Fig 3B) and overall conjunctive strength (Fig 4B) analyses (both FDR p<.05). The supplement (Table S1 and Fig S4) details the 5 subcortical ROIs selected, with accompanying network information from the CAB-NP brain parcellation. Identifying subcortical ROIs ensured sufficient signal-to-noise for the analyses of representational dynamics. These criteria also informed the decision to confine cortical analyses in Fig 5-8 to the average for each of the 12 CAB-NP networks.

Note that the cortical network averages excluded all subcortical regions affiliated for each network, to ensure clean separation between cortical and subcortical contributions. The averages across cortical networks and subcortical regions were also included to aid timecourse visualization (versus presenting 19 timecourses in a single plot) in Figs 5-8. Note that the ventromedial multimodal (VMM) network was included in the subcortical average, given that the bilateral perirhinal regions within this network are intimately linked to the function of subcortical MTL regions (O’Reilly et al., 2014; Tomás Pereira et al., 2016).

To interrogate the association between pattern separation (neural task similarity) and rule strength (Fig 7), Spearman’s correlation was computed between the block-wise neural task similarity timecourse and the rule strength timecourses (3 compositional and 1 conjunctive, each averaged across the 4 practiced tasks). The resulting subject-level rho values were submitted to group random effects statistics, at both the regional and network-averaged levels.

We also extended estimation of the compositional and conjunctive representations into practice task blocks presented in the second Test session. Each of the 4 practiced tasks were repeated a further 15 times in this session. The rule templates were once again calculated by averaging over relevant held-out novel tasks in the Test session, to prevent circularity. Visual impressions of the resulting rule strength timecourses were formalized by averaging the estimates over 6-block epochs spanning early Practice (blocks 1-6), late Practice (blocks 31-36), early Test (blocks 37-42) and late Test (blocks 46-51).

We followed similar analytic conventions in the analyses that estimated compositional and conjunctive rule strength separately for the task switching sub-conditions of the practiced tasks (2 Switch tasks versus 2 NoSwitch tasks, see earlier Methods for design details, see Fig 8C-D for results). Compositional and conjunctive rule strength was estimated across Practice session blocks per the approach in Fig 4A, separately for each of the 4 tasks, with the resulting timecourses averaged into Switch and NoSwitch conditions before subsequent analysis. These timecourses were presented visually in Fig S9B, and also summarized in the text as the epoch-averaged rule strength (early i.e. block 1-6, or late i.e. block 31-36), as above. The construction of the brain-behavior index linking cortical conjunctions to task switching behavioral costs is detailed in the later “Behavioral analyses” section.

### Dynamic trajectory analysis

For both the neural task similarity and rule strength analyses, trends over time in the observed block-to-block timecourses were quantified using a simple linear regression approach (Mill et al., 2016):

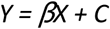

where Y is the neural measure (neural task similarity or compositional/conjunctive rule strength), X is time (block 1-36), C is a constant, and ꞵ is a coefficient capturing the linear trajectory over time (positive=increasing, negative=decreasing). Robust regression was again used to ensure improved resilience of the estimated coefficients to noise/outliers in the timecourses (Holland & Welsch, 1977). Dynamic trajectories were estimated for each individual subjects’ neural timecourses, and then submitted to group random effects statistics to assess significance.

### Behavioral analyses

Standalone analyses of behavior focused on performance accuracy (% correct) and reaction time (RT), averaged across the 3 trials presented within a task block. We plotted behavioral timecourses across chronologically presented blocks for accuracy (Fig 2C) and RT (Fig 2D) to verify recovery of standard learning effects (improved performance with practice). For analyses that contrasted behavior against chance performance accuracy (Fig 2B and Fig 2C), even though chance was technically 25% (given that 4 button responses were available, 2 for each hand), we considered 50% (2 button options on a given hand) as chance under the assumption that subjects usually correctly identified which hand to use for a given task.

Analyses of the fMRI data that tested for brain-behavior associations computed Spearman’s rank correlation between the relevant blockwise imaging measure (neural task similarity or rule representation strength), separately with behavioral accuracy and behavioral RT. Associations were generally stronger with RT, which is why we focused on this behavioral measure in the Results. For the neural task similarity measure (Fig S3), this was computed between the neural task similarity timecourse (one timecourse computed across all tasks, see above) and the cross-task averaged RT timecourse. For rule strength (Fig 6), this was computed separately between a given rule type’s timecourse (sensory, logic, motor or conjunctive) and RT for a given task. The resulting Spearman’s timecourses were then averaged across the 4 practiced tasks prior to random effects group statistics.

For the task switching analyses, we firstly calculated behavioral costs of switching, by contrasting RT (Fig 8C) and accuracy (Fig S9A) between Switch versus NoSwitch tasks. Trends in the behavioral timecourses were summarized via epoch averages that followed previous conventions (early: block 1-6, late: block 31-36). We also computed a brain-behavior association index separately for each cortical network and subcortical region (Fig 8D). This linked blockwise differences in conjunctive rule strength between task switching conditions to blockwise differences in behavior (RT) between the same conditions. For each subject, Spearman’s correlation was computed between rule strength (NoSwitch minus Switch) and behavioral RT (NoSwitch minus Switch), such that negative rho values captured an index of behavioral switch costs in the appropriate direction (i.e. quicker RT for NoSwitch versus Switch). These subject-level rho values were then analyzed via a random effects approach at the group level (detailed in the last section).

We selected Spearman’s correlation as our method of brain-behavioral association given its established sensitivity to monotonic relationships that may be linear or non-linear (e.g. exponential/power law relationships). To validate this choice, we conducted stepwise regression modeling of behavioral RT to examine whether linear and/or non-linear temporal relationships were observable in the behavioral data. This analysis predicted subject-level RT timecourses from linear and non-linear (power law) variables, fit in separate models, and simultaneously in a combined model. The coefficient of determination (R^2^) was used to compare model accuracy across these variations (linear only, non-linear only and combined). The exponent of the non-linear predictor was estimated flexibly for each subject by first fitting a linear regression to log-transformed linear time (X) and RT (Y) variables.

We also conducted a control analysis involving partial Spearman’s correlation, which regressed out the linear time variable prior to computing the various brain-behavioral associations (accompanying Fig S3, Fig 6 and Fig 8D). Considering that we had already removed the linear trend and various technical and physiological artifacts via expanded nuisance regression, this additional conservative step was an attempt to exhaustively rule out any residual temporal artifacts in the brain-behavior associations.

### Cross-region association of rule representation dynamics

For each subject, we computed cross-region associations between the rule representation timeseries (estimated over Practice session blocks, per Fig 4A). Our chosen association measure was graphical LASSO (GLASSO), which is a multivariate method based on partial correlation that is capable of identifying direct associations from indirect and confounded associations. It also uses L1 regularization to prevent overfitting to noise. GLASSO was shown recently to yield improved inter-session reliability and validity compared to other univariate and multivariate methods, in a study focused on optimizing brain functional connectivity methods (Peterson et al., 2023). Whilst that study computed associations amongst “first-level” fMRI BOLD timeseries, we anticipated similar benefits when computing associations amongst “second-level” rule representation timecourses. In this way, our approach can be considered a variation of previous informational connectivity approaches (Coutanche & Thompson-Schill, 2013).

Each GLASSO model fit rule strength timeseries for all cortical networks and subcortical regions simultaneously. Separate GLASSO models were run for the various combinations of rule types: compositional-compositional, compositional-conjunctive and conjunctive-conjunctive. Each yielded a 17-by-17 adjacency matrix of cross-region association estimates (partial correlation r values), which was symmetric across the diagonals. Each off-diagonal matrix element captured cross-region associations of rule representation dynamics, between cortical networks and subcortical regions (defined in Fig S4). For the compositional-compositional analysis, we estimated the associations separately for each rule type (sensory-sensory, logic-logic and motor-motor) and then averaged them before computing group-level statistics. Similarly, for the compositional-conjunctive analysis, we estimated the associations separately for each compositional rule type (sensory-conjunction, logic-conjunction, motor-conjunction) and then averaged them. Averaging over compositional rule types was supported in both cases by the results being very similar for the separate rules. The resulting adjacency matrices were visualized after group averaging (Fig 8E, Fig S11), and summarized by taking the average of off-diagonal elements for each cortical network/subcortical region (i.e. each column in the subject’s matrix; Fig 8F, Fig S11). Note that whilst the compositional-compositional and conjunctive-conjunctive analyses targeted propagation of the same type of compositional rule representation across regions, the compositional-conjunctive analysis targeted transformation (binding) of rule representations across regions, through associating two different rule representation types (compositions and conjunctions). The compositional-conjunctive analysis was also unique in having a meaningful diagonal, which captured within-region associations between the two rule types (Fig S11F). Whilst the directness and regularization features of the adopted GLASSO approach raise clear advantages over other methods (e.g. Pearson correlation, multiple linear regression), it is worth noting that it lacks clear directionality of influence (contrasting with e.g. Granger Causality), which is a limitation to address in future work.

### Statistical analysis

All presented statistics – neural task similarity rho values, rule representation strength betas, behavioral association rho values, and rho values for associations of neural task similarity and rule strength – used 2-sided one-sample t-tests against 0 to assess significance, treating subject-level estimates as a random effect. Where specified in the Results, paired sample t-tests (2-sided) were used for contrasts across conditions. Spearman rho values underwent the Fisher-z transformation to improve normality prior to running the t-tests. T-statistics from these contrasts are depicted in all figures, thresholded using the false discovery rate (FDR p<.05; Benjamini & Hochberg, 1995) procedure to correct for multiple comparisons across regions, networks or block timepoints (where appropriate). The Supplementary Table lists significant regions for all regional analyses, with accompanying t and p values. This includes the CAB-NP region number (1-360 are cortical, and 361-718 are subcortical), allowing for extraction of anatomical information with reference to CAB-NP documentation (https://github.com/ColeLab/ColeAnticevicNetPartition).

## Acknowledgements

The authors acknowledge support by the US National Institutes of Health under award R01 MH109520, and support by the US National Science Foundation under award 2219323. The authors acknowledge the Office of Advanced Research Computing (OARC) at Rutgers, The State University of New Jersey, for providing access to the Amarel cluster and associated research computing resources that have contributed to the reported results. We would also like to thank Julia Hamilton, Emily Winfield, Doug Schultz, Richard Chen, Nicole Lalta, Kirsten Petersen and Kaustubh Kulkarni for assistance with piloting and data collection.

## Data availability

Preprocessed fMRI data for each subject will be uploaded to an Open Science Foundation repository (with the access link provided) upon publication.

## Code availability

Custom MATLAB code used to empirically quantify the strength of rule type (i.e. compositional and conjunctive) representations on a block-to-block basis in the practiced task data is provided here https://github.com/ColeLab/CPRO2_learning_release.

## Supplementary Information

### Task information was reliably decodable in the new C-PRO2 paradigm

We used a multivariate classification approach to verify that neural representations evoked by our C-PRO2 paradigm contained decodable task information. This served as a sanity check demonstrating that manipulated task features were decodable in fMRI multivariate activation patterns, which preceded later analyses testing for specific formats of the multivariate representational geometry (i.e. contributions of compositional and conjunctive rule representations) over learning.

4-way classification of practiced task identity was conducted using a minimum distance algorithm (Haxby, 2001; Mill et al., 2020; Mur et al., 2009; Spronk et al., 2020) across Practice session blocks. As a reminder, the C-PRO2 paradigm permutes 64 tasks (each composed of 3 rules, spanning sensory, logic and motor rule types), of which 4 tasks were assigned to the repeated practice condition, and the remaining 60 to the novel condition (performed only once). This analysis focused on decoding the identity of the 4 practiced tasks from regional multivariate fMRI activation patterns (chance decoding = 1/4 tasks = 25%).

Templates for each practiced task were estimated (trained) in the held-out Test phase via a general linear model, which included regressors for each of the 4 practiced tasks and 60 regressors of no interest for the novel ones. For a to-be-classified practice task block, a “correct” decision was output if the Spearman correlation was higher with the correct (matching) Test session task template, compared to correlations with any of the other incorrect templates. This was repeated for each block and each region (using within-region voxels/vertices as features) to create regional task information timecourses. The cross-block averages for each subject were then subjected to group-level averaging and random effects statistical thresholding (one-sample t-test for classification accuracy versus chance 25%, FDR p<.05).

The results in Fig S1 reveal a number of regions containing decodable task information, encompassing cortex (e.g. motor, higher-order sensory and lateral PFC regions) and subcortex (including bilateral cerebellum, bilateral dorsal striatum and left thalamus). The Supplementary Table lists all significant regions for this analysis and accompanying statistics. Network averaging of the regional decoding accuracies (right panel Fig S1) revealed broad significance across networks (10/12 were significantly above chance, FDR p<.05), albeit with relative gradation between networks. This spatially distributed-but-graded task information profile has been observed across multiple neural data modalities (Cole et al., 2016; Kauvar et al., 2020; Mill et al., 2022; Siegel et al., 2015). The relatively stronger decodability seen in motor, cognitive control and higher sensory network regions is consistent with prior multi-task studies (Cocuzza et al., 2020; Cole et al., 2013; Ito et al., 2017; Mill et al., 2022). Strong decodability of the cerebellum is also consistent with prior reports linking it to flexible multi-task cognition (King et al., 2019; Nakai & Nishimoto, 2022).

These results confirmed that multivariate activation representations evoked by our new paradigm contained decodable task information, with a spatial profile consistent with prior multi-task findings. Note that unlike the neural task similarity (Fig 3) and rule representation strength (Fig 5) results, dynamic trajectories estimated from the task information timecourses were relatively stable over practice (i.e. no region had a significant positive or negative trajectory). Hence, while these task information results provided a sanity check of the functional relevance of multivariate activations evoked by our task, our analyses focusing on dynamics of representational geometry (neural task similarity and rule representation strength) in the main manuscript were more sensitive in elucidating the neural basis of task learning.

**Fig S1.**
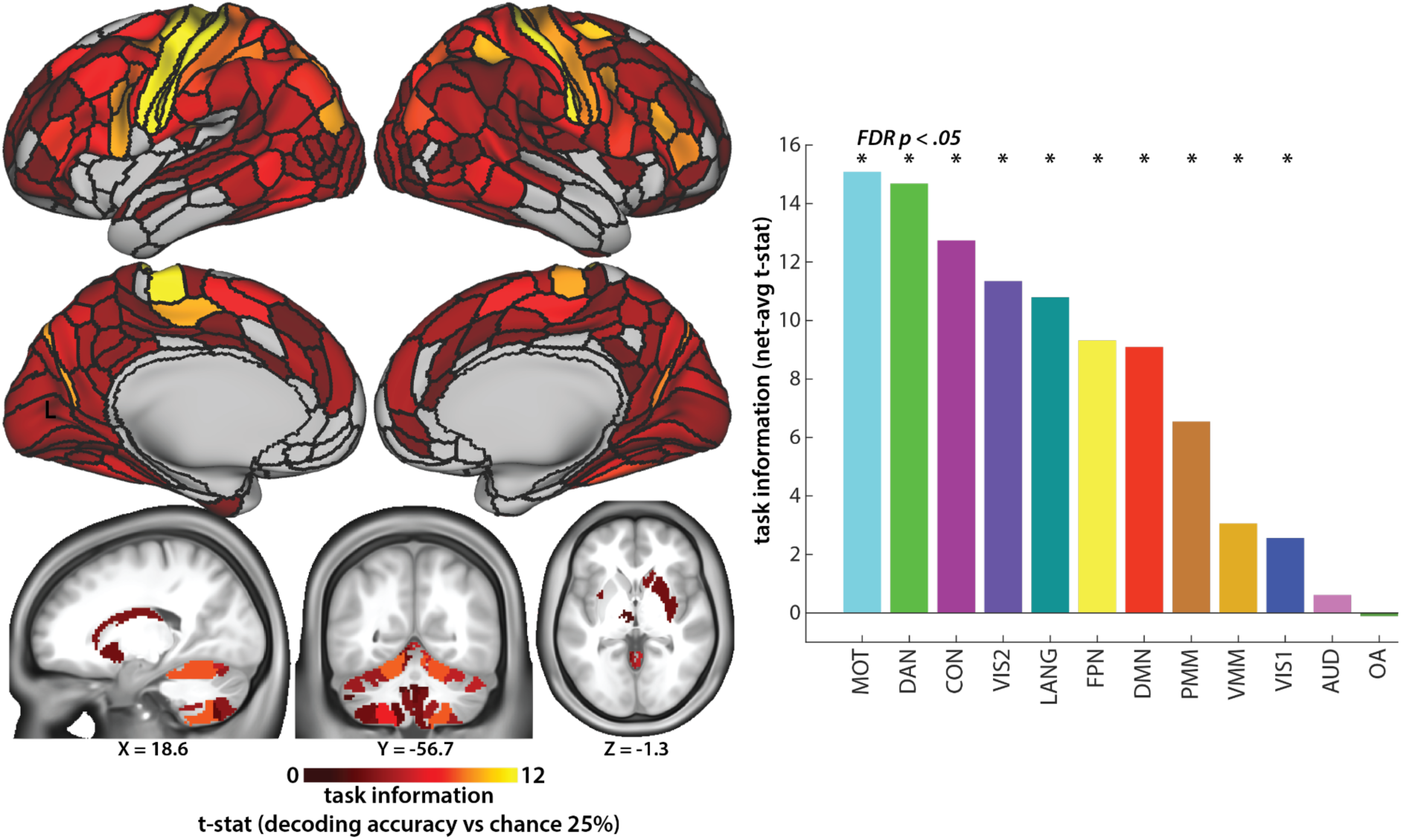
Multivariate classification reveals reliable decodability of neural task information, with a distributed-but-graded spatial profile. Results of the 4-way classification of practiced task identity in the Practice session (see text for details), at the regional level (left panel) and network-averaged level (right level). Task information was reliably and broadly decodable, spanning established cortical and subcortical regions. The network summaries revealed broad significance across networks (10/12), but with relatively higher task information decodability in motor, cognitive control and higher sensory networks.

### Whole-brain pattern separation results

Fig S2 depicts the results of applying the neural task similarity analysis at the whole-brain spatial scale (treating all 718 cortical/subcortical regions as features). This was done to depict an exemplar neural similarity timecourse which was the basis of estimating dynamic trajectories in the regional analyses in Fig 3, as well as to probe whether pattern separation processes emerged at this more macroscopic scale. The results revealed a significant negative trajectory for neural task similarity consistent with pattern separation (Fig S2A), and this held when the 12 individual C-PRO2 rule types were visualized separately (Fig S2B). Correlating the whole-brain neural task similarity timecourse with behavioral accuracy and RT (analogous to the regional results in Fig S3) again revealed beneficial associations with behavior: accuracy increased as neural similarity decreased, t(43)=-3.37, p=.002; RT decreased as neural similarity decreased: t(43)=6.71, p<.001. The findings suggest that pattern separation learning principles hold at macroscopic spatial scales and across multiple task rule types.

**Fig S2.**
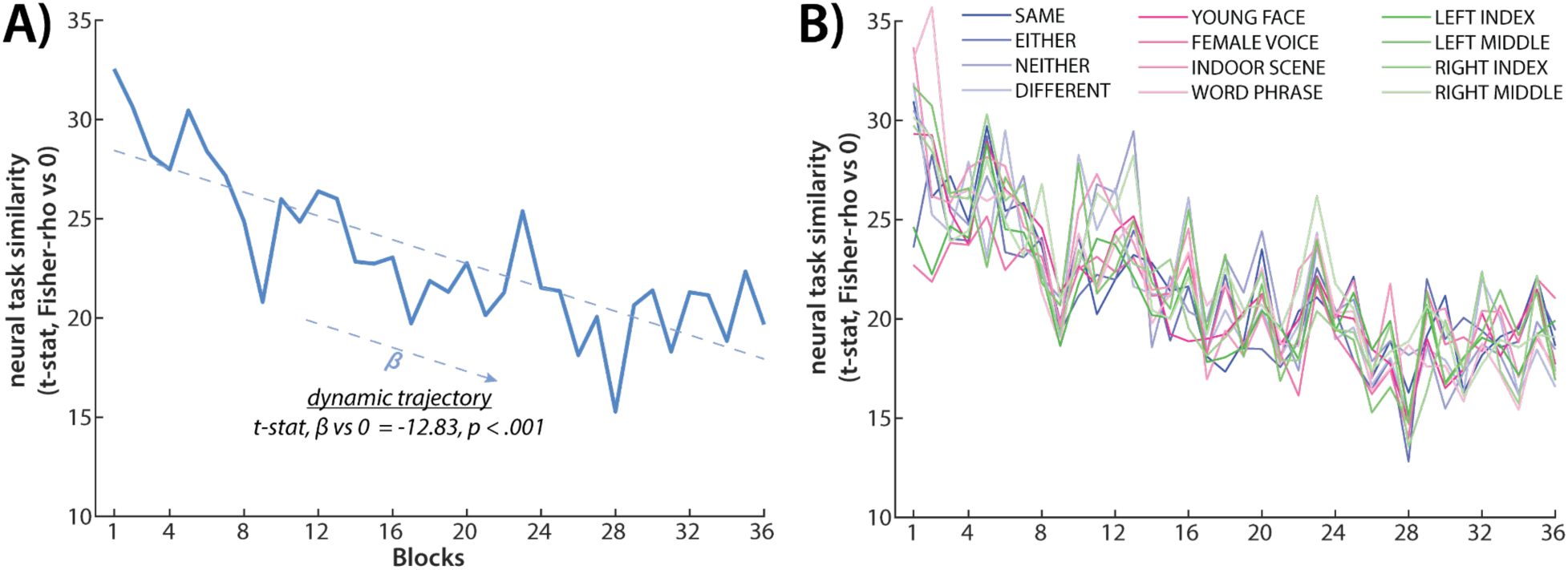
Whole-brain neural task similarity decreases consistently across C-PRO2 rule types. **A)** Neural task similarity computed at the whole-brain scale (all 718 CAB-NP regions as features) reveals a negative trend consistent with pattern separation. Linear line of best fit and results of dynamic trajectory analysis are overlaid. **B)** Neural task similarity visualized separately for each of the 12 C-PRO2 rules (clustered in the legend by logic, sensory and motor rule types). Rule-specific timecourses were computed by averaging pairwise similarity values involving each rule within subjects, which yielded non-independent timecourses due to the one-to-one rule-to-practiced task assignment (see Methods). Although this precluded statistical comparison across rules, the visualized group-averaged timecourses highlight the generality of the negative trajectory (pattern separation) over time.

### Increase in pattern separation over repeated practice is associated with behavioral improvement

Brain-behavior associations involving the cognitive control networks survived a partial correlation approach that regressed out time prior to computing the associations, in an exhaustive attempt to remove temporal confounds/artifacts (see Methods). Surviving accuracy associations: FPN t=-2.74, p=.009; CON t=-3.98, p<.001; DAN t=-2.31, p=.026. Surviving RT associations: FPN t=2.01, p=.050; CON t=2.21, p=.033.

**Fig S3.**
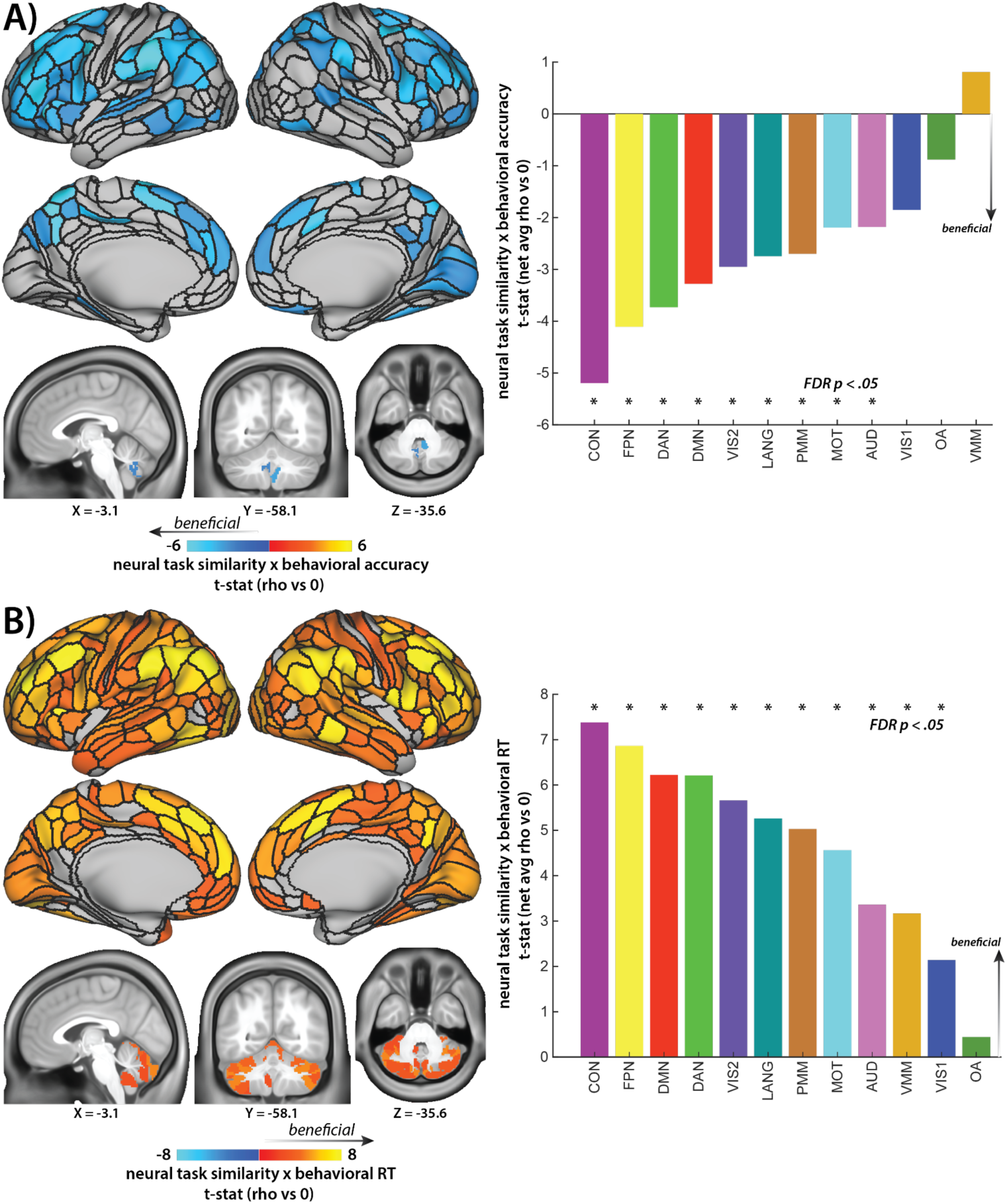
Degree of neural task similarity decrease (pattern separation) correlates with behavioral improvement with practice. **A)** Associations between regional neural task similarity and behavioral accuracy (% correct), computed via Spearman’s correlation (one-sample t-stat for Fisher-rho vs 0, at FDR p<.05), for practiced tasks in the Practice session. Negative values reflect beneficial associations i.e. as neural task similarity decreased over learning, behavioral accuracy increased. This was observed in multiple cortical regions and two subcortical regions in bilateral cerebellum. **B)** Equivalent analysis for RT, with positive values reflecting beneficial associations i.e. as neural task similarity decreased over learning, RT decreased. Multiple cortical regions reached significance, as did multiple subcortical regions (all in bilateral cerebellum). Network averages of the regional associations are presented to the right in both panels, revealing prominent loci for these behavioral influences in the CCNs, DMN and higher sensory networks.

### Selection criteria and information for subcortical regions-of-interest included in the rule strength analyses (Figs 5-8)

**Table S1.** Subcortical regions of interest. Selection was based on strong pattern separation and conjunctive strength effects, formalized as regional significance in both the neural task similarity (one-sample t-test for similarity trajectory beta versus 0, Fig 3B) and overall conjunctive rule strength (one-sample t-test for rule beta coefficient versus 0, Fig 4B) analyses, both thresholded at FDR p<.05.

| CAB-NP region index | CAB-NP network region label | ROI label used in Figs 5-8 |
| --- | --- | --- |
| 433 | Visual2-10_L-Cerebellum | L/R hemispheres averaged together to form VIS2 cerebellum region (cVIS2) |
| 438 | Visual2-15_R-Cerebellum |  |
| 495 | Cingulo-Opercular-15_L-Cerebellum | L/R averaged together to form CON cerebellum region (cCON) |
| 501 | Cingulo-Opercular-21_R-Cerebellum |  |
| 529 | Dorsal-Attention-10_L-Cerebellum | DAN cerebellum region (cDAN) |
| 584 | Frontoparietal-28_R-Cerebellum | FPN cerebellum region (cFPN) |
| 659 | Default-23_L-Hippocampus | L/R averaged together to form DMN hippocampus region (hippo) |
| 661 | Default-25_R-Hippocampus |  |

**Fig S4.**
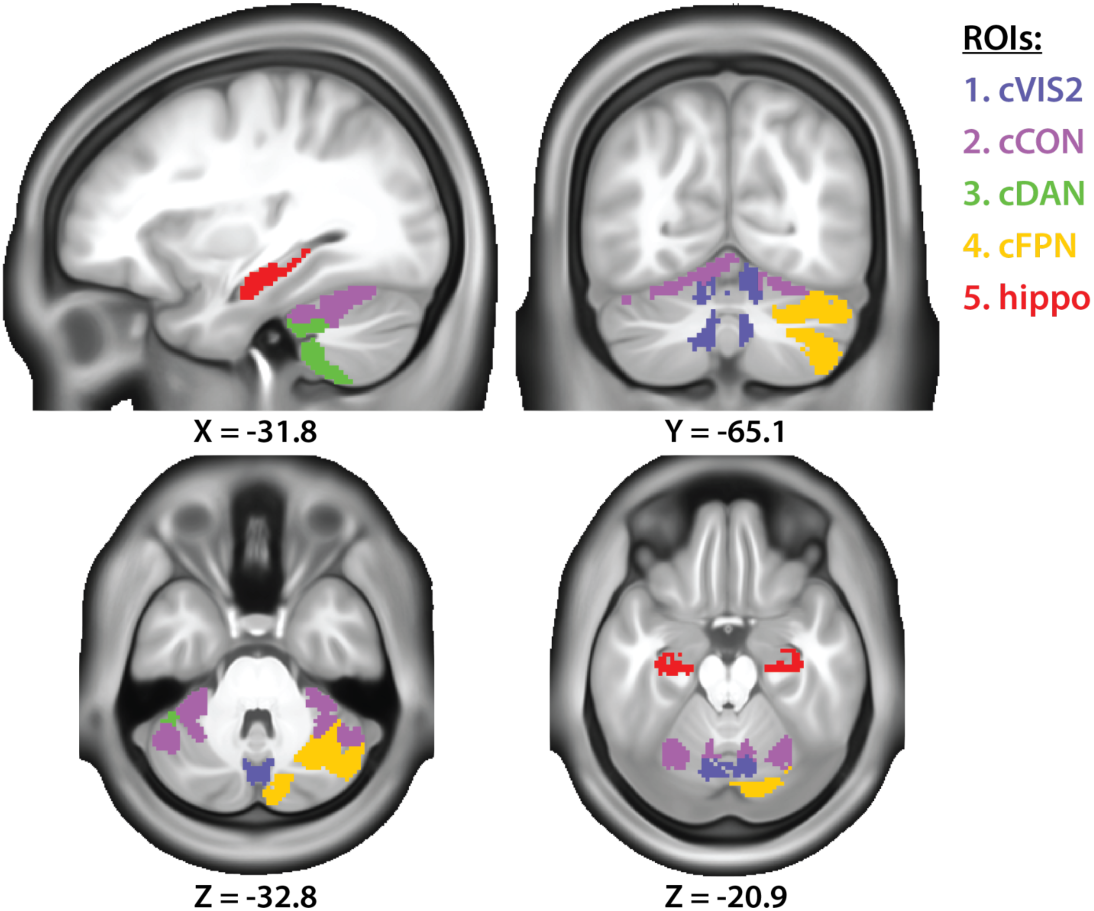
Visualization of subcortical ROIs used for rule representation strength analyses. See Table S1 for anatomical and functional information for these 5 regions, referencing the CAB-NP network parcellation. ROI colors denote CAB-NP network affiliation.

### Expanded interpretation (with a ground truth simulation) of early negative cortical conjunction estimates as reflecting compositional interference

In the dynamic rule strength analyses presented in Figure 5A, cortical conjunctive strength was numerically negative early in the course of learning, before becoming increasingly positive over Practice (Figure 5B) and Test (Figure 8A-B) sessions. Whereas our focus in the main analysis was on the statistically robust positive trajectory of the cortical conjunctions, here we expand on the correct interpretation of the early negative conjunctive estimates. To preview, the early negative cortical conjunction estimates do not reflect some kind of “negative code” for strong engagement of conjunctions, but rather reflect representational interference arising from strong early engagement of the compositional representations.

From a statistical standpoint, it is worth highlighting that the conjunction predictor in our analysis (Fig 4A) is the multiplicative interaction of the 3 compositional main effects (sensory, logic and motor rule templates). Hence, we can draw on prior statistical discussions of how to correctly interpret the interaction term in multiple linear regression models (Jaccard et al., 1990). Firstly, interpreting the sign of an interaction benefits from referencing the sign of the main effects. In this case, the early compositional estimates (main effects) were all positive (Fig 5B-D). When main effects are positive, a negative interaction (as observed early in cortex; Fig 5A) indexes a suppressive effect, wherein strong engagement of all main effects yields a reduced (subadditive) effect on the outcome variable than anticipated by summing those effects. In the present context, strong early engagement of the 3 compositional rule templates yielded a suppressive effect on the observed regional activation patterns (our outcome variable), which can be considered a form of representational interference.

Note that this interpretation aligns closely with the schematized rule representation dynamics in Fig 1C. Early in task learning, cortical compositional representations are strongly engaged, driving increasing representational overlap (interference) between tasks. As this early compositional engagement progressively weakens, this leaves the task-relevant conjunction (yellow patch in Fig 1C) to dominate, as indexed by increasingly positive conjunctive estimates. As highlighted in Fig 1C, one computational benefit of increasingly relying on the conjunctions is reduced representational interference, as the engagement of the more general compositional representations may trigger engagement of task-irrelevant representations.

We operationalized the above principles in an expository simulation (Fig S5) with a known ground truth, which closely followed the ideas laid out in Fig 1C. We began by creating rule representation templates for three compositional rules (comp1-3 in Fig S5) and their multiplicative interaction (conj), in a 6 vertex-by-6 vertex cortical region. We then simulated observed activation patterns over time (outcome variables Y), which followed the ground truth principle of strong early compositional engagement that progressively weakens towards relying on the conjunctive region patch. This process spanned an initial distributed activation pattern (YT1) and a final sparser, conjunctive-recruiting activation pattern (YT10). Blockwise activation patterns were sampled between these start and end patterns, via simple linear spacing of the vertex-wise activations over 10 blocks. Note that the number of sampled blocks is arbitrary with respect to actual learning effects in the brain.

Generally, this simulation aimed to elucidate fundamental statistical principles governing how variation in the sign of an interaction term (conjunction) emerges from concurrent variation in the constituent main effects (compositions).

**Fig S5.**
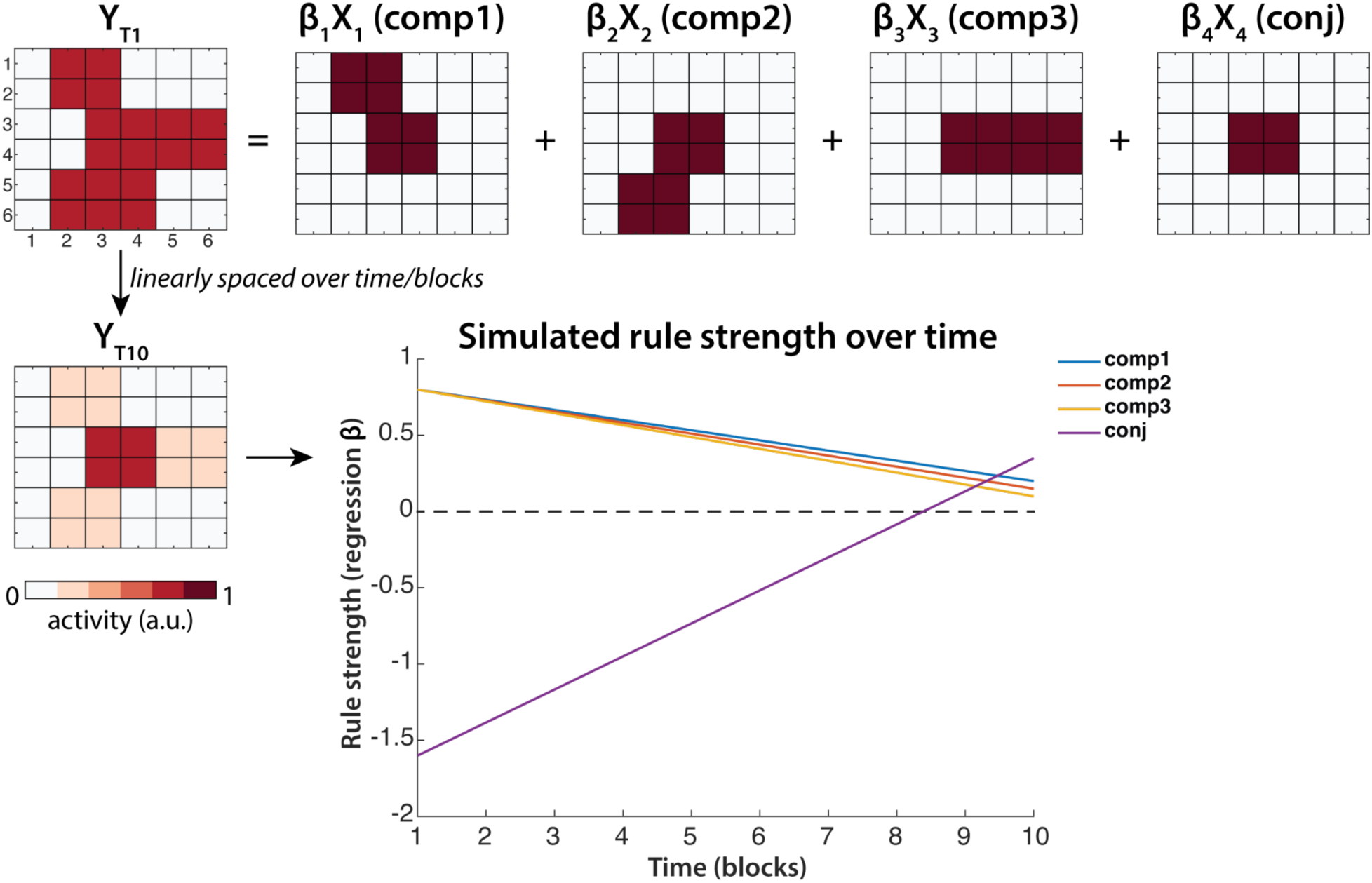
Ground truth simulation capturing progression of conjunctive rule strength estimates over time, from early negative to increasingly positive. See text for details.

The results of fitting the rule templates to these simulated block-to-block activation patterns are depicted in the lower right plot of Fig S5. As predicted, the multiple linear regressions fit to the simulated data recovered early negative conjunctive betas when compositions were strongly engaged. These became increasingly positive over time, as the conjunctions became more strongly and selectively engaged relative to the larger compositions. The pattern of simulated results therefore closely parallels our statistical explanation above, and our empirical estimates of cortical conjunctive strength across Practice (Fig 5A) and Test (Fig 8A) sessions. This suggests that our interaction-based estimate does provide a continuous index of how strongly conjunctions are engaged over time in the brain.

### Recovery of equivalent learning dynamics via an alternative method of estimating compositional representations

To demonstrate generalizability of the observed learning dynamics across variation in how rule representations were estimated, we applied an alternative, established method of detecting compositional representations: cross-condition generalization performance (CCGP; Bernardi et al., 2020). This is related to standard multivariate decoding approaches (e.g. train a decoder to differentiate between conditions A vs B over a mix of sub-conditions C and D, then test decoding of A vs B on held-out data that is also a mix of C and D), but is more systematic in training/testing across sub-conditions (e.g. train for A vs B on C, test for A vs B on D, and vice versa, then average the decoding results to yield a measure of CCGP).

To clarify, with our original approach, to estimate a compositional template for the SAME rule in a practiced task SAME + YOUNG FACE + LEFT INDEX, we averaged regional activations over all instances of SAME in the held-out novel tasks (15 total). With our CCGP implementation, we instead iterated this template-matching procedure over specific combinations of SAME and other rules. For example, a “correct” template would be selected as the regional activation for the novel task activation SAME + YOUNG FACE + LEFT MIDDLE, with 3 “incorrect” templates selected by swapping out SAME for the other 3 logic rules (i.e. DIFFERENT + YOUNG FACE + LEFT MIDDLE, EITHER + YOUNG FACE + LEFT MIDDLE, NEITHER + YOUNG FACE + LEFT MIDDLE). Rule strength would then be computed as the similarity (Spearman correlation) of the to-be-decoded practiced task regional activation with the correct compared to incorrect rule templates (i.e. correct minus incorrect similarity computed separately for all 3 incorrect templates and then averaged). This CCGP implementation therefore used a 4-way minimum distance classification approach (Mill et al., 2020; Mur et al., 2009; Spronk et al., 2020). The process was repeated for all other presentations of SAME in the novel tasks, each time computing “incorrect” templates by swapping out SAME with the three logic rules, with the resulting decoding values averaged to give a CCGP estimate of logic rule compositional strength on that practiced task block. Repeating this for all practiced task blocks in the Practice session, and for all 3 rule types (logic, sensory and motor) yielded compositional rule strength timecourses for each region, as with our original approach (Fig 4 and Fig 5).

Note that our original approach to detecting compositional rule representations was also CCGP-esque in that the testing observations (which were averaged over to create templates) were each separate novel task conditions by design, rather than random/unstructured condition repetitions. Hence, our original approach also permits inferences of compositionality and abstraction over and above regular decoding and representational similarity approaches. Nevertheless, CCGP is a more exhaustive alternative, albeit one that trades off this exhaustiveness with greater noise in the representational templates due to not averaging over multiple novel task conditions to create them.

Fig S6A plots the cross-block averaged compositional rule strength for each of the rule types. This again recovered sensible positive region peaks i.e. higher-order visual regions for sensory rules, lateral prefrontal and parietal regions for logic rules, and somatomotor regions for motor rules (see Supplementary Table for accompanying statistics). As before, compositional rule strength was overall higher in cortex compared to subcortex (cross-block and cross-rule averaged rule strength: cortical t = 12.11, subcortical t = 6.58, both p<.001; cortical > subcortical rule strength, t=9.68, p<.001). Fig S6B plots the dynamic trajectories for the rule strength timecourses for cortical networks and subcortical regions, again revealing weakening compositional rule strength for sensory and motor rule types. Fig S6C presents associations between compositional rule strength and behavioral RT, again finding associations generally in the harmful direction, particularly for sensory and motor rules. Dynamic trajectory and behavioral association effects were comparatively weaker for the logic rule type, which is a trend also present in the original analyses (Fig 5C and Fig 6C). More targeted comparison of effects between rule types is beyond the scope of the present report, but is planned for future work.

**Fig S6.**
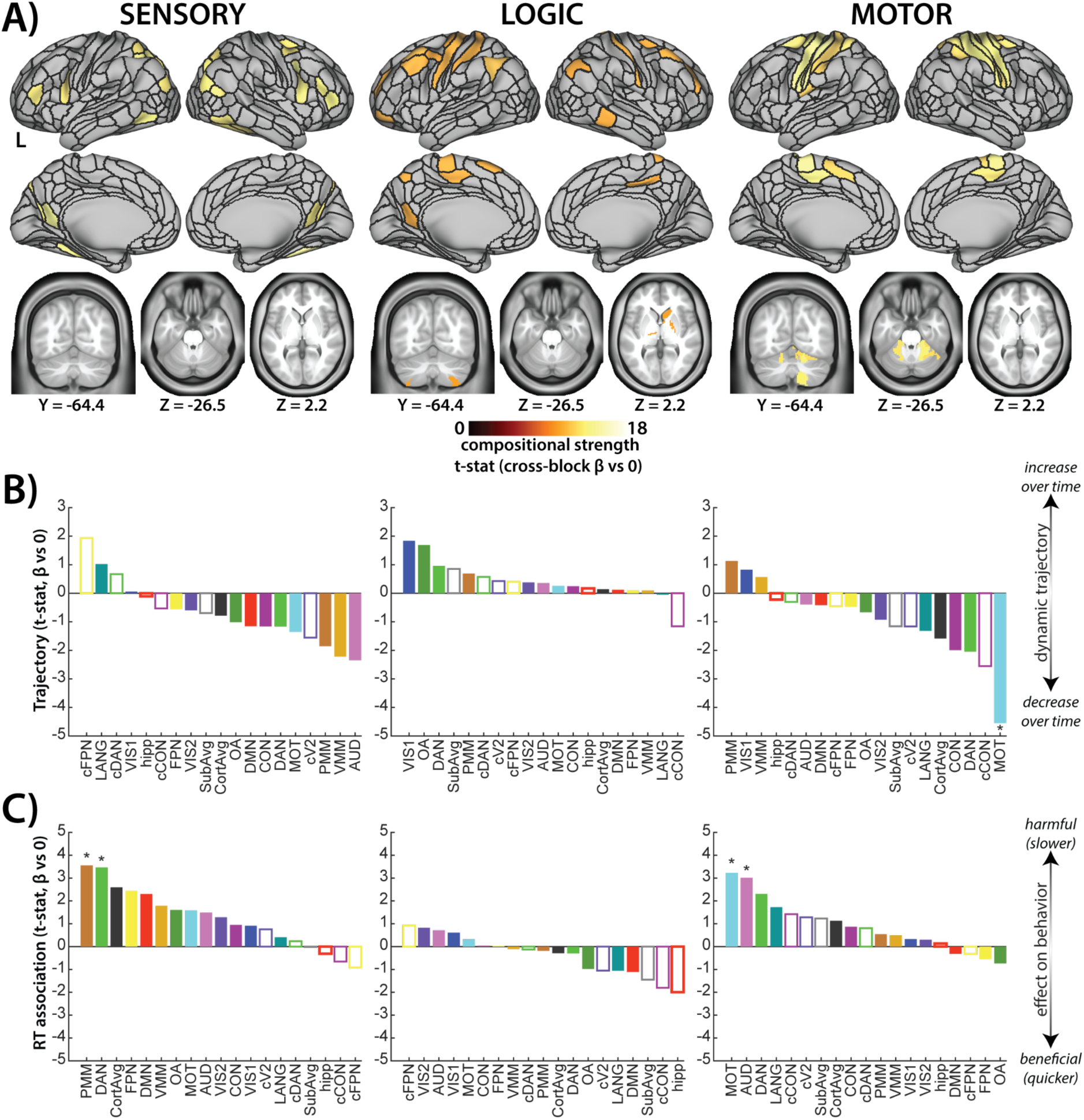
CCGP implementation reveals similar dynamics of compositional representation over learning. Columns plot the CCGP results for sensory, logic and motor rule types. **A)** Cross-block compositional rule strength revealed sensible positive peaks, and overall stronger compositional representations in cortex compared to subcortex (top 5% positive regions plotted; see Supplementary Table for statistics after top 5% and FDR thresholding). **B)** Dynamic trajectory analysis after averaging regional effects into cortical networks and subcortical regions (per Fig S4). This revealed general weakening (negative trajectories) of compositional rule strength over practice. **C)** Brain-behavior associations: weakening of compositional rule strength over time was associated with improved behavior (quicker RT). Asterisks in panels B-C denote significance at FDR p<.05.

Overall, the CCGP results in Fig S6 strongly accord with the compositional learning dynamics uncovered by our original approach, highlighting the robustness of the reported effects to analytic variation. The pattern of CCGP findings was virtually identical when using alternative similarity metrics than Spearman correlation (Pearson correlation and cosine similarity).

Note that our examinations of generalizability were unable to include the only alternative method of estimating conjunctive representations (that we are aware of; Kikumoto & Mayr, 2020), due to inherent features of our task design. To clarify, implementing the approach of Kikumoto and Mayr would require fitting template representational similarity matrices (RSMs), capturing task-to-task similarity embedded in the study design, to actual RSMs, capturing task-to-task similarity of a region’s evoked activation patterns. The diagonal of the actual RSM codes for similarity in repetitions of the same task, and is critical to allow estimation of conjunctive representations. In our case, populating the diagonal for novel tasks in the actual RSMs was precluded due to the lack of repetition of the novel tasks, which were each only presented once to preserve their novelty. This highlights an inevitable trade-off in multi-task fMRI designs, between presenting a more diverse array of tasks versus repeating those individual tasks, whilst keeping session durations per subject within reasonable limits. Nevertheless, modifying the PRO task design to enable direct comparisons of our approach to estimating conjunctive representations with that of Kikumoto & Mayr is planned for future work.

### Compositional rule strength in the second Test session: evidence for re-engagement after a lapse in task performance (linked to increased cognitive demand)

The results of estimating compositional rule strength in the second Test session are provided in Fig S7, with cortical/subcortical average timecourses in the left panels, and epoch averages (over 6 blocks as in Fig 8B) in the right panels. The pattern was consistent across all 3 rule types: compositions were strongly engaged early in the first Practice session (when tasks were novel) and subsequently weakened within that session, before transiently being re-engaged at the start of the second Test session and then again numerically weakening. These impressions were formalized by contrasting epoch estimates over time, separately for each rule type type, and separately for cortex and subcortex. Blue lines in Fig S7 highlight significant rule strength differences between the epochs (via paired t-test contrasts, FDR p<.05).

All 3 rule types, across cortex and subcortex, showed a significant increase in compositional rule strength for the early Test epoch, compared to the early Practice and late Practice epochs (all FDR p<.05). Compositional rule strength also numerically declined within-session: for Practice (early Prac > late Prac for all rule types, across cortex/subcortex, reaching significance for Motor rules in cortex, t=2.62, p=.012, FDR-corrected) and numerically (albeit non-significant) for Test (early Test > late Test for all rule types, reaching a trend level of non-significance for Motor rules across cortex, p=.119, and subcortex, p=.122). Note that contrasts involving the late Practice and late Test epochs are hard to interpret, given the difference in overall practice duration for Practice (36 blocks) and Test (15 blocks) sessions. That is, compositional strength in the late Test epoch may have returned to a comparably low level as late Practice if more practice had been provided at Test.

Whilst the above pattern was consistent across cortex and subcortex, cortical compositions were overall more strongly engaged (via paired-sample t-test contrasts at each epoch, cortex > subcortex, all FDR p <.05).

**Fig S7.**
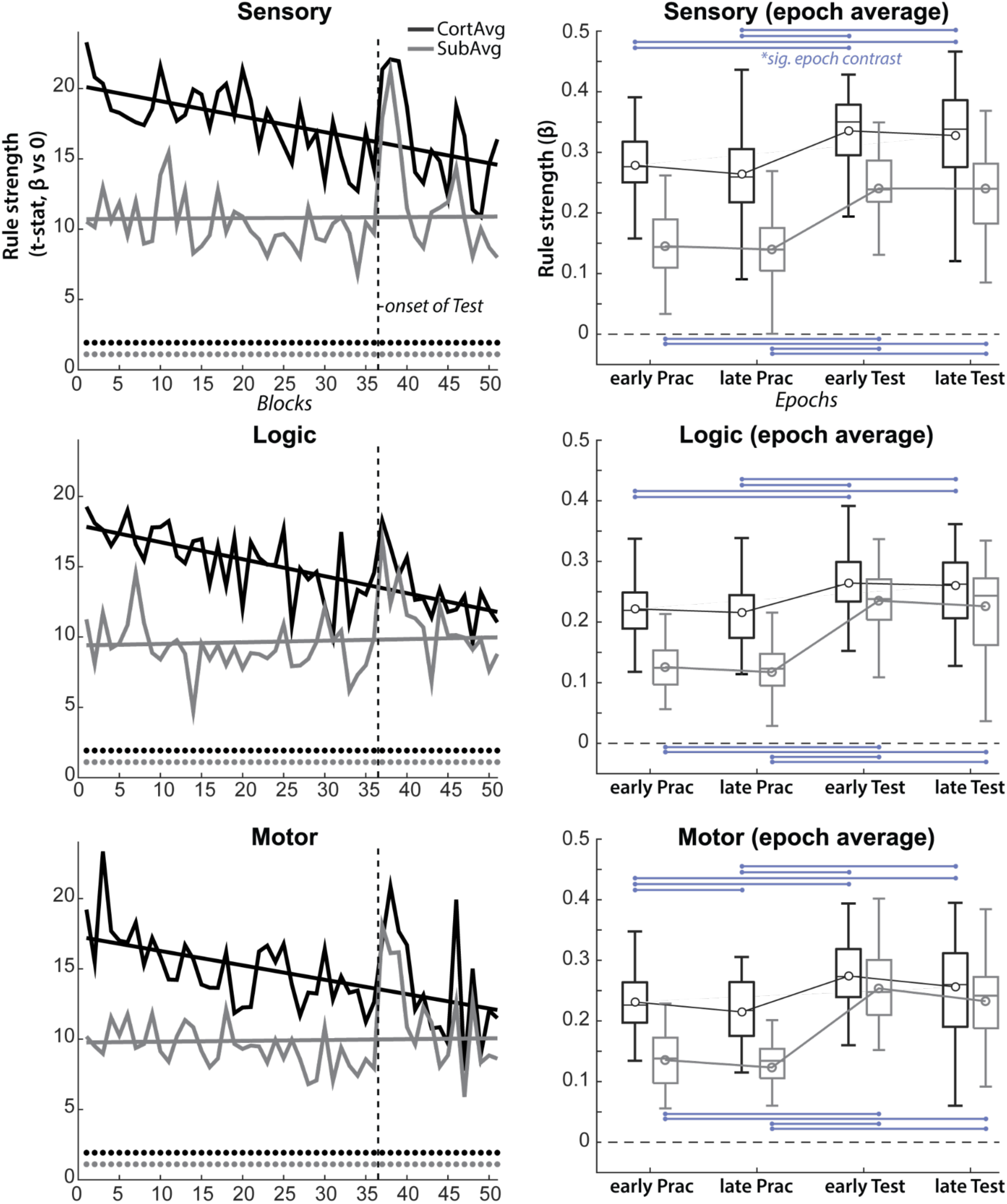
Dynamics of compositional representation across the first Practice and second Test sessions. Plotting conventions are the same as Figure 8A-B.

Hence, the Test session analyses again reveal a differing dynamic profile for compositional versus conjunctive (Fig 8A-B) rule strength, which lends support to our hypothesis that they capture separate key computations in cognitive task learning. However, the Test session results extend and add nuance to the role of rule compositions. Specifically, the increase in compositional strength for early Test suggests that even practiced tasks can elicit strong compositional engagement, which we interpret as reflecting an increase in cognitive demand due to the 1-7 day lapse in task performance. This link to increased demand can accommodate both the strong compositional engagement observed when tasks were novel (early Practice, Fig 5), as well as the strong re-engagement after the performance lapse (early Test, Fig S7). In both cases, the initial compositional engagement was followed by within-session weakening, and parallel strengthening of cortical conjunctions. Note our use of “parallel” here – if compositions and cortical conjunctions were mutually orthogonal, the early Test compositional increase would not have been observed at the same time as cortical conjunctions were strongly evoked. The observed parallel engagement of compositions and conjunctions at the start of the Test session therefore may align with prior work recovering similar parallelized representational geometries, allowing a balance between abstraction and specialization (Bernardi et al., 2020).

The link between compositional engagement and cognitive demand was strengthened by behavioral analyses of the Test session in Fig S8. The figure contrasts behavior in the last block of the Practice session with the first practiced task block in the Test session (i.e. the first block after the lapse in performing the practiced tasks). Behavioral performance worsened after the lapse in performance, for both accuracy (Prac last block mean = 86.5%, Test first block mean = 74.6%, t=2.56, p=.014) and RT (Prac last block mean = 2421.6 ms, Test first block mean = 2750.2 ms, t=3.96, p<.001). This supports the notion that cognitive demand increased at the start of the Test session, and our theory that this was compensated for by the concurrent increase in compositional engagement (Fig S7). The overall pattern suggests that compositions are more generally engaged under conditions of high demand, which require elevated cognitive control. This is in opposition to cortical conjunctions which more continuously track the extent/robustness of practice across varying demands (i.e. there was no comparable ‘peak’ for cortical conjunctions in the early Test epoch).

**Fig S8.**
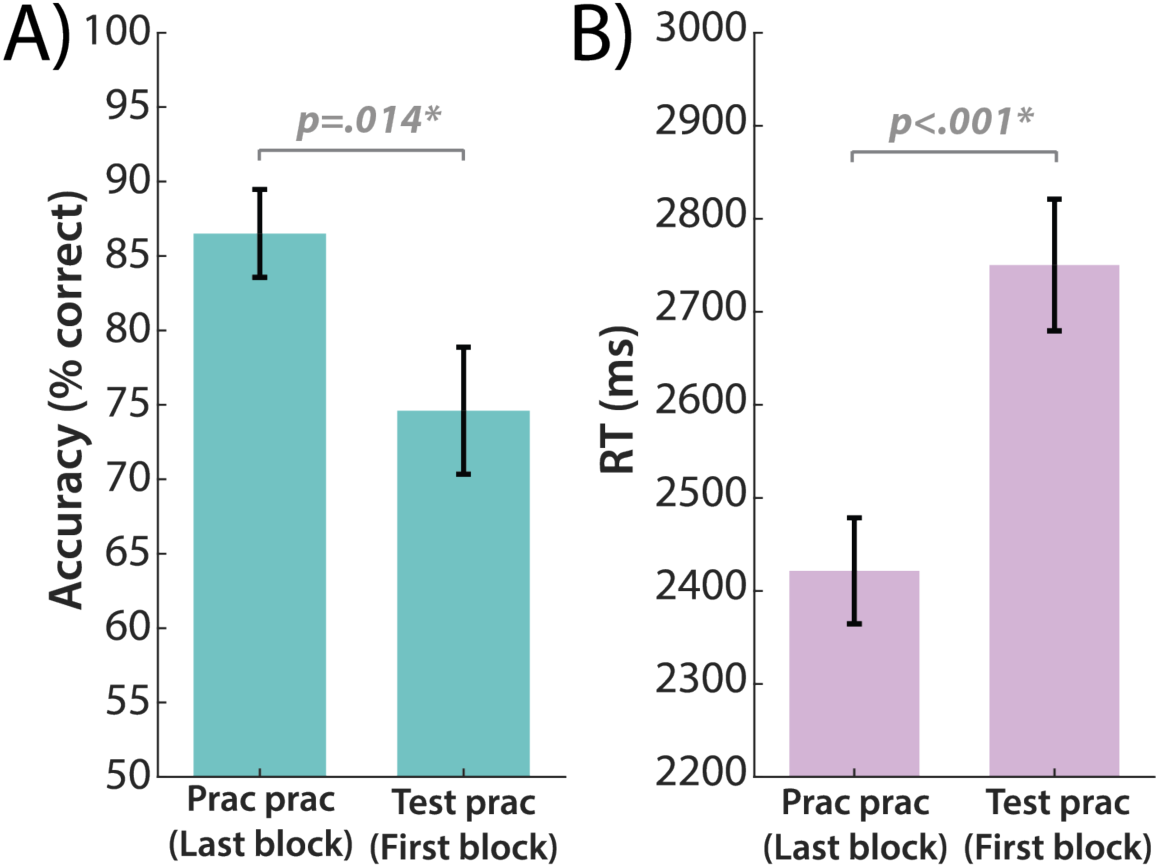
Behavior worsened (cognitive demand increased) after a lapse in practiced task performance. Both panels compare behavior between the last practiced task block of the Practice session with the first practiced task block of the Test session. **A)** Results for behavioral accuracy. **B)** Results for behavioral reaction time (RT). P values correspond to paired t-test contrasts.

### Rule strength estimated separately for Switch and NoSwitch practiced task conditions

**Fig S9.**
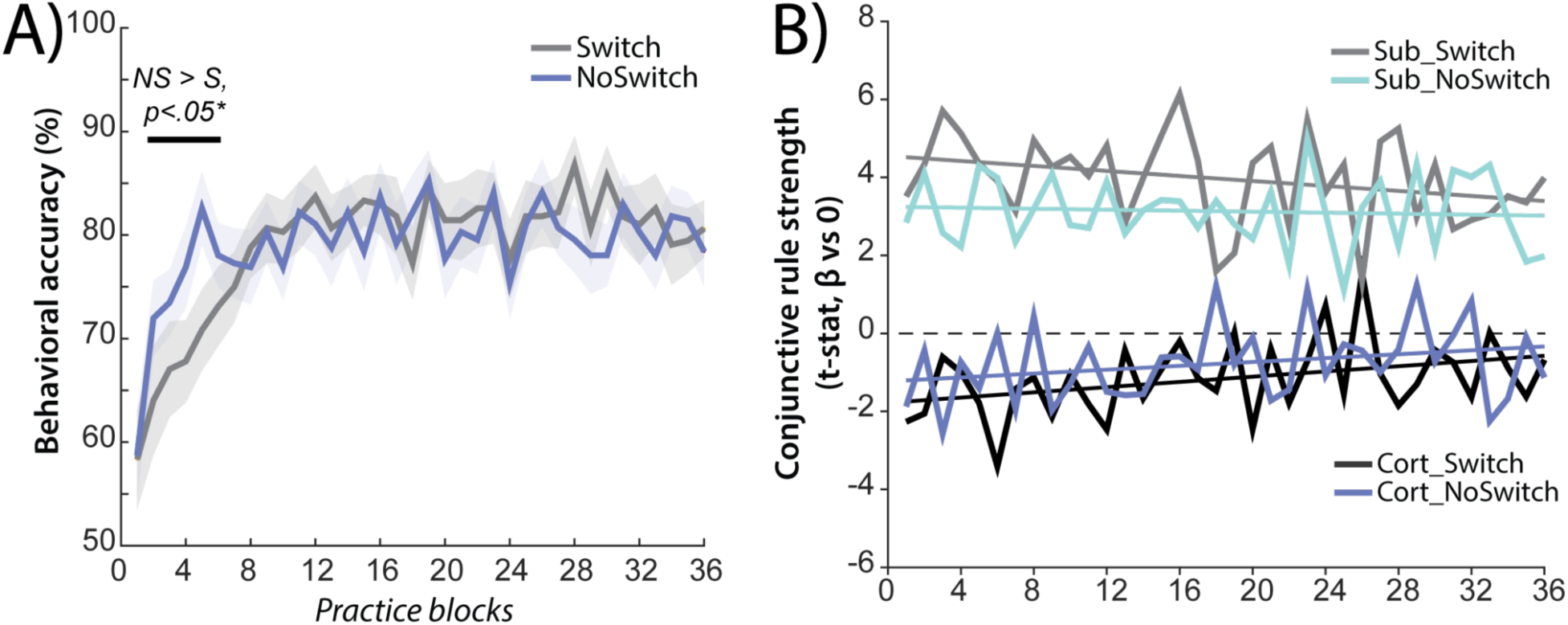
Analyses of conjunctive rule strength across task switching conditions. **A)** Behavioral cost of task switching: Switch practiced tasks yielded lower accuracy than NoSwitch practiced tasks early in the session (block 1-6). **B)** Conjunctive rule strength during the Practice Session, estimated separately for Switch/NoSwitch tasks, and cortical networks/subcortical regions. See main Results text for details.

**Fig S10.**
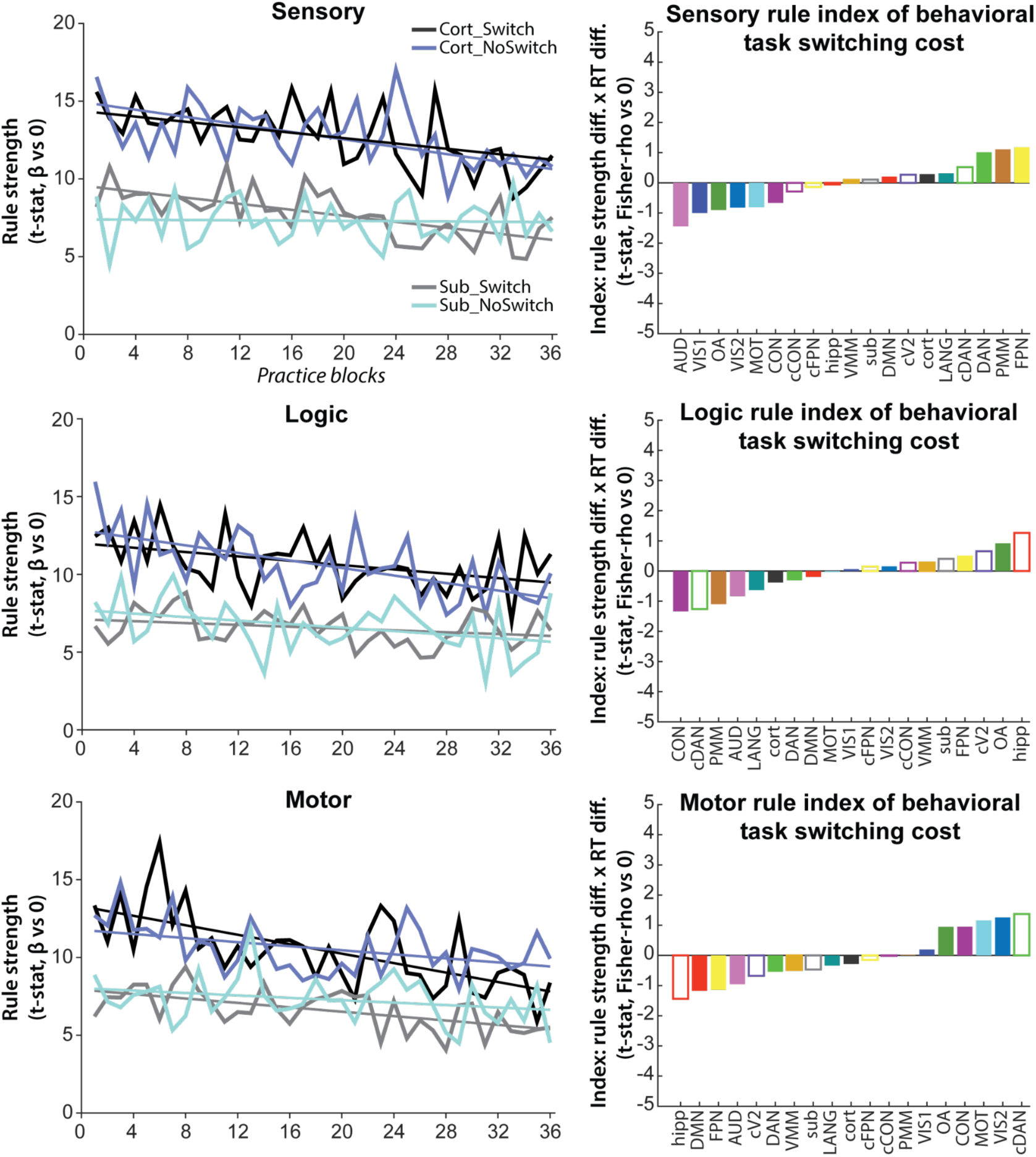
Analyses of compositional rule strength across task switching conditions. Rows plot the results for the 3 compositional rule types (sensory, logic, motor). Left columns plot compositional rule strength timecourses, separately for Switch and NoSwitch practiced task conditions, separately for cortex and subcortex. This reveals similar trends as in earlier analyses that collapsed across these conditions: compositional rule strength is overall greater in cortex compared to subcortex, and generally decreases with practice. Differences in session-level (block averaged) compositional rule strength across Switch vs NoSwitch conditions were overall modest, across sensory (cortical, Switch > NoSwitch, t=1.09, p=.281; subcortical Switch > NoSwitch, t=1.78, p=.082), logic (cortical, Switch > NoSwitch, t=0.29, p=.795; subcortical, Switch > NoSwitch, t=0.00, p=.997), and motor rules (cortical, Switch > NoSwitch, t=0.00, p=.999; subcortical NoSwitch > Switch, t=-0.10, p=.992). Right panels plot brain-behavior associations between compositional rule strength differences (NoSwitch minus Switch) and behavioral RT differences (NoSwitch minus Switch). None of the compositional rule types, across cortical networks and subcortical regions, yielded a reliable association with behavior. This further highlights the uniqueness with which FPN conjunctive rule strength indexes task switching costs, as reported in the main manuscript (Fig 8D).

### Cross-region associations of rule representation dynamics

**Fig S11.**
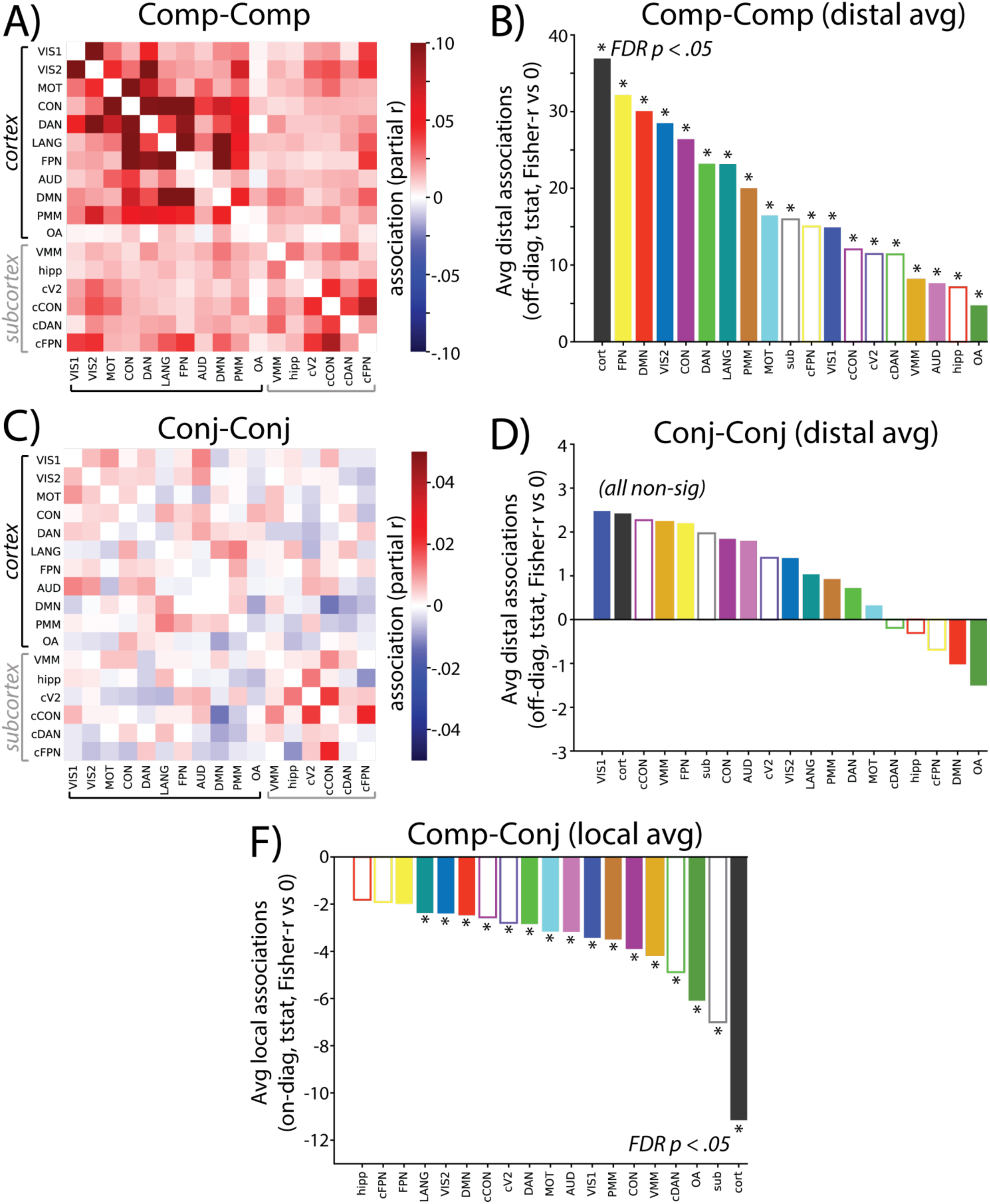
Cross-region association results. **A)** Group-averaged matrix for compositional-compositional rule dynamics and **B)** the off-diagonal (distal) matrix average. Panels **C)** and **D)** plot the equivalent for conjunctive-conjunctive dynamics. **F)** The on-diagonal (local) average for compositional-conjunctive rule associations. Note that this is the only matrix on-diagonal that was meaningful, due to capturing local (within-region) associations between different rule types (compositional and conjunctive) over time. The generally negative values (following the group matrix in Fig 8E) served to recapitulate the opposing within-region trajectories of compositional and conjunctive representations in Fig 5. Refer to the main manuscript for further interpretation.

